# A key region of molecular specificity orchestrates unique ephrin-B1 utilization by Cedar virus

**DOI:** 10.1101/724138

**Authors:** Rhys Pryce, Kristopher Azarm, Ilona Rissanen, Karl Harlos, Thomas A. Bowden, Benhur Lee

## Abstract

The prototypic henipaviruses (HNV), Hendra (HeV) and Nipah (NiV), are emergent zoonotic pathogens responsible for frequent and fatal outbreaks of severe disease in domestic animals and humans. The HNV attachment glycoprotein (G) is a critical determinant of host-species and cell-type tropism. Utilization of highly conserved B-type ephrin ligands as functional entry receptors engenders HNVs with a broad permissive host range, accounts for zoonotic spillover, and is closely aligned with observed disease pathologies. Recent studies have uncovered numerous divergent clades of HNVs globally. Cedar virus (CedV), the closest relative of HeV and NiV identified to date, can establish experimental infections, yet has not been observed to cause overt disease. While the apathogenic phenotype may be attributed to a lack of P-gene derived interferon antagonists, the V and W accessory proteins, additional determinants of differential HNV pathobiology could be involved. Here, through comparative functional and structural analysis of CedV-G, we characterize molecular interactions critical to viral entry. We demonstrate that CedV possesses a unique cellular entry receptor repertoire which, in addition to functional utilization of the common HNV receptor, ephrin-B2, includes the hitherto uncharacterized interaction with ephrin-B1. Crystal structures reveal a conserved recognition mode between diverse HNV-G proteins and their distinct ephrin receptors and identify a region of molecular specificity within CedV-G that is a key determinant of ephrin selectivity. This work provides a platform for understanding the functional diversity and varied receptor tropism characteristics of HNV glycoproteins that will facilitate assessment of the pathogenic potential and transmissibility of newly discovered and uncharacterized HNVs.

## Introduction

The prototypic henipaviruses (HNV), Hendra (HeV) and Nipah (NiV), are biosafety level four (BSL4) pathogens responsible for severe human disease that is associated with rapid onset and case fatality rates that can exceed 90% [1-3]. Combined, the extreme disease pathologies, absence of a licensed vaccine, paucity of medical intervention options, and zoonotic potential delineate HNVs as an acute and persistent threat to global biosecurity, economy, and health [4]. Whilst zoonotic spillover is typically associated with transmission from chiropteran reservoirs, or via infection of domestic animal intermediates, such as pigs and horses, transmission is not restricted to cross-species spillover events. Direct human-to-human spread is frequent and highlights the pandemic potential of HNVs [5-7].

Serological studies suggest that HNVs occupy a broad geographic range coincident with, but not restricted to, the home range of reservoir bat species of the order *Chiroptera* [8, 9]. Whilst there is evidence for the existence and spillover of previously uncharacterized HNVs in Africa and Central- and South-America [10-15], an accurate appraisal of the human impact of such HNVs is likely hindered by the spectrum of clinical outcomes inherent to diverse HNV species [16]. Indeed, the indirect association of a novel HNV, Mójiāng virus (MojV), with the death of three miners in China highlights the potential of non-chiropteran hosts as reservoirs of lethal HNVs [17]. The putatively rat-borne MojV utilizes an ephrin-independent host-cell entry pathway and possesses a structurally distinct attachment glycoprotein [18]. The continued discovery and emergence of novel HNV species underscores the indeterminate global health threat that they pose [4, 12].

Cedar virus (CedV) is a *Henipavirus* species isolated from the excreta of *Pteropus* bat colonies in Queensland, Australia [19]. Although geographically, genetically, and serologically related to the highly virulent prototypic HNVs, CedV is apathogenic in small animal models [19]. The stark disparity in HNV pathogenesis has been attributed, in part, to the lack of an otherwise conserved RNA editing site and the alternate reading frame coding capacity for accessory proteins within the CedV phosphoprotein (P) gene [19]. In NiV and HeV, RNA editing facilitates the production of the accessory proteins, V and W, which are capable of antagonizing the interferon (IFN) response. The absence of these immunomodulatory accessory proteins in CedV results in a failure to mitigate the anti-viral effects of the type I IFN response and likely represents a critical factor in determining infection outcomes [20].

The single-stranded negative-sense RNA genome of HNVs encodes two surface glycoproteins: the receptor-binding glycoprotein (G) and the type I viral fusion protein (F), which work in concert to orchestrate cellular entry [21-23]. Binding of the HNV-G to cell-surface receptors belonging to the ephrin ligand family initiates pH-independent activation of F, triggering a fusion cascade that results in the ultimate merger of viral and cellular membranes. HNV-G proteins comprise a short N-terminal cytosolic region, single-pass transmembrane domain, oligomerization-mediating and fusion-activating stalk region, and C-terminal receptor-binding β-propeller domain [24, 25]. Orthologs of the two established HNV receptors, ephrin-B2 and ephrin-B3, are extremely well conserved across numerous reservoir and vector species, and are recognized by all ephrin-tropic HNV-G proteins with a conserved binding mode [26-28]. Utilization of ephrins as cellular entry receptors is fundamental to the broad cell-type and species tropism of HNVs and underscores key features of HNV zoonosis and pathogenesis [29, 30].

Despite lacking canonical type I IFN antagonists (V and W) [19, 20], CedV does possess a functional C accessory protein, the counterparts of which exhibit type I IFN antagonism in NiV [31-34]. Furthermore, CedV is able to establish a productive, albeit self-limiting, infection in Syrian hamsters that is more robust when inoculated via the intranasal versus intraperitoneal route [20]. Together, these observations suggest that the identity of cellular receptors for CedV and the efficiency with which they are utilized may constitute additional modifiers of pathogenicity. Here, we sought to delineate the functional entry receptor repertoire of CedV and elucidate the molecular determinants of receptor specificity. In our integrated structural and functional analysis, we demonstrate that in addition to utilizing the common HNV receptor, ephrin-B2, CedV utilizes ephrin-B1, a receptor with no precedent of HNV usage. Structural analyses reveal that whilst CedV-G conforms to a universal mode of HNV-G mediated ephrin receptor engagement, subtle structural features of the glycoprotein contribute to its unique ephrin ligand specificity. These data highlight functional diversity amongst HNV-G proteins and provide mechanistic insight into potential modulators of HNV pathobiology.

## Results

### The crystal structure of CedV-G reveals a conserved receptor-binding architecture

Although reported to utilize the common entry receptor, ephrin-B2 [19, 35], CedV-G is genetically distinct from all characterized ephrin-tropic HNVs (26%, 28%, and 31% identical to GhV-G [Ghana virus], HeV-G, and NiV-G, respectively) [27]. To assess the extent to which this sequence divergence is reflected at a structural level, we determined the crystal structure of the CedV-G receptor binding domain to 2.78-Å resolution. Two essentially identical molecules of CedV-G were present within the crystallographic asymmetric unit (root-mean-square deviation [RMSD] 0.3 Å across 416 equivalent Cα atoms), with the only region of notable structural variation localizing to the β6-S2–S3 loop (Fig. 1a). Electron density permitted modelling of the entire receptor-binding domain of CedV-G included in the crystallized construct (residues K209–C622) (Fig. 1b).

**Figure 1:**
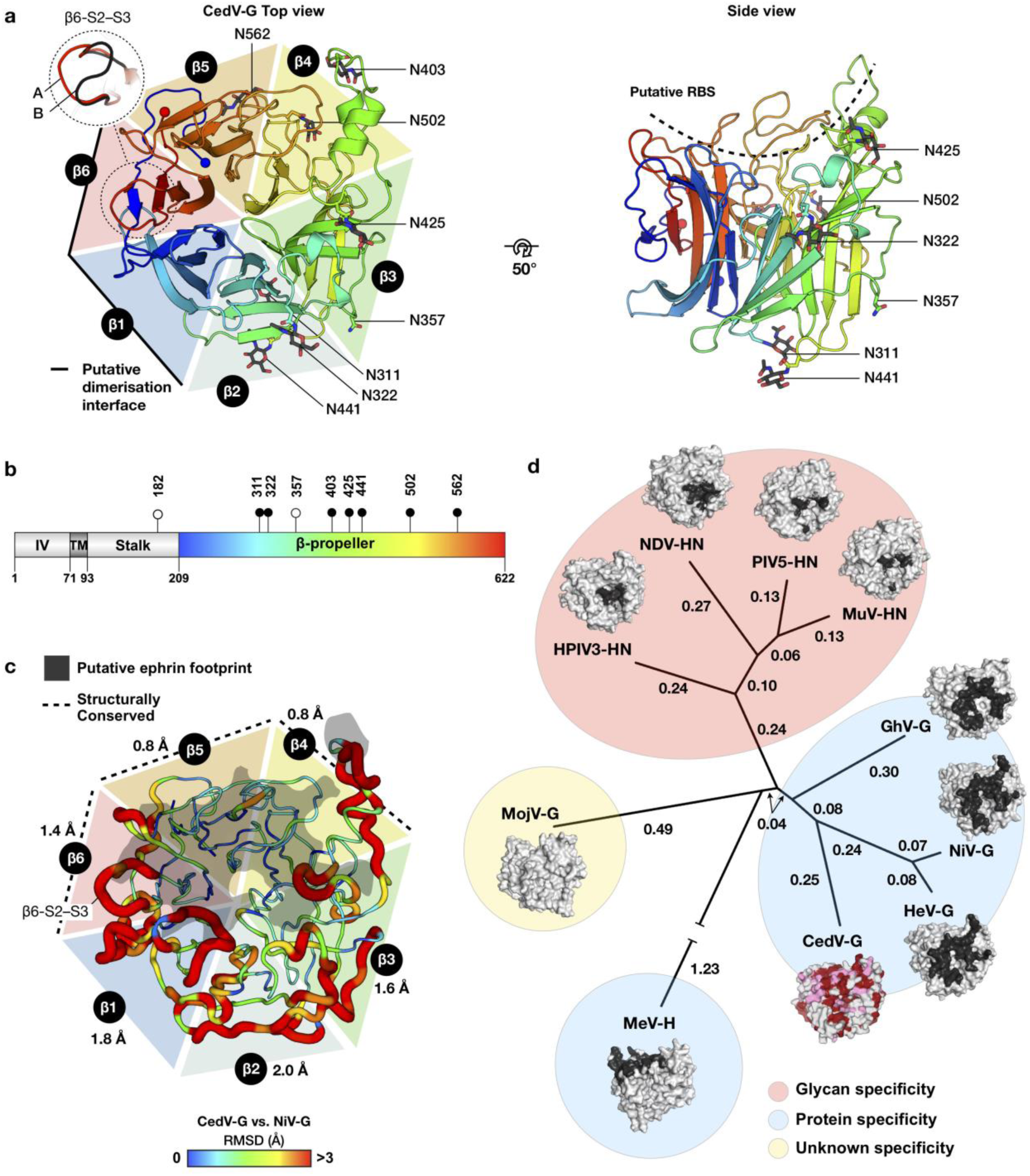
The CedV-G β-propeller exhibits regions that are structurally conserved with ephrintropic HNV-Gs suggestive of a preserved mode of receptor recognition. **a**, The structure of CedV-G is displayed as a cartoon colored from N- to C-terminus (blue to red), with the termini shown as spheres. The approximate extent of each of the six ‘blades’ of the β-propeller is delineated by a colored triangle, labelled β1–β6. The region of highest variability (β6-S2–S3 loop) between the asymmetric unit copies (molecule ‘A’ and ‘B’) of the protein is shown with an inset panel (dashed line). Molecule A is shown in rainbow and the β6-S2–S3 loop of molecule B is shown in black. The protein is displayed in a ‘top’ view (left) and rotated 50° to reveal a side view on which the putative receptor binding site (RBS) is depicted with a dashed line. N-linked glycans are shown as sticks and colored according to constituent elements. Asparagine residues from all eight N-linked glycosylation sequons are displayed as sticks. The putative dimerization interface contributed by the β1 and β6 blades is denoted with a solid black line. **b**, Domain organization and salient features of CedV-G. The attachment-mediating β-propeller domain, transmembrane (TM), and intra-virion (IV) regions are labelled. Putative N-linked glycosylation sites are displayed as pins, with sites occupied in the crystal structure colored black. The extent of the crystallized construct is colored as a rainbow as in **a. c**, Structural comparison of CedV-G and unliganded NiV-G (PDB accession code: 2VWD). Due the high level of structural similarity between NiV-G and HeV-G [45], an HeV-G comparison is omitted for clarity. Root-mean-square deviation (RMSD) between aligned Cα residues is depicted by both color (in a gradient from blue to red with increased RMSD) and tube width (thin to thick with increased RMSD). Residues that failed to align or exhibited RMSDs greater than 3 Å were assigned values of 3 Å. The average RMSD across each blade is displayed next to the respective blade label and the more structurally conserved region of the molecule (β4–β6) is indicated with a dashed line. The NiV-G–ephrin-B2 interface is displayed as a grey shadow superposed onto the structure of CedV-G **d**, Structure-based phylogenetic analysis of paramyxovirus receptor binding proteins places CedV-G amongst ephrin-utilizing HNV-G proteins. Pairwise distance matrices were calculated with SHP [76] and plotted with PHYLIP [77], using the structures of unliganded receptor-binding glycoproteins, where available. The corresponding structures are shown in surface representation with previously characterized receptor binding surfaces shown in dark grey. The structure of CedV-G is colored according to sequence conservation with NiV-G, identical residues are red and similar residues are pink. Measles virus hemagglutinin (MeV-H) branch is truncated for illustrative purposes. Structures utilized for the analysis were: the Ghanaian bat henipavirus G (GhV-G; PDB 4UF7), Nipah virus G (NiV-G; PDB 2VWD), Hendra virus G (HeV-G; PDB 2X9M), CedV-G, MeV-H (PDB 2RKC), Mòjiāng virus G (MojV-G; PDB 5NOP), human parainfluenza virus 3 hemagglutinin-neuraminidase (HPIV3-HN; PDB 1V3B), Newcastle disease virus HN (NDV-HN; PDB 1E8T), parainfluenza virus 5 HN (PIV5-HN; PDB 4JF7), mumps virus HN (MuV-HN; PDB 5B2C). Branches of the resultant tree are labelled with the calculated evolutionary distances.

The receptor-binding domain of CedV-G adopts the canonical six-bladed β-propeller fold utilized by ephrin-tropic (NiV-G, HeV-G, and GhV-G) [26, 36-38], and ephrin-independent (MojV-G) HNVs [18]. Each of the six blades (β1–β6) comprise four antiparallel β-strands that assemble in a toroidal arrangement and form a central depression at the membrane-distal ‘top’ surface of the molecule (Fig. 1a). Like the prototypic HNVs, the β2 and β3 blades are decorated with extended loops that form three short 3_10_ helical segments (η1–η3) and three α-helices (α1–α3). Eight disulphide bonds stabilize the fold, seven of which are conserved amongst other HNV-G proteins (Supplementary Fig. 1), indicative of structural importance in stabilizing the β-propeller. The additional disulphide linkage in CedV-G (C310–C376) links blades β2 and β3 at the base of the molecule.

The receptor-binding domain of CedV-G possesses eight N-linked glycosylation sequons, three more than the prototypic HNV-Gs. Electron density corresponding to an asparagine-linked N-acetylglucosamine moiety was observed at seven sites (Fig. 1a & b and Supplementary Fig. 2a). The distribution of glycan sites within CedV-G is consistent with previous studies that implicate the glycan-free β1 and β6 blades in the formation of an homo-dimeric interface as part of the higher-order assembly of virion-displayed HNV-Gs [37, 39, 40]. Only one of these sequons, N502^CedV^, is conserved with HeV-G (N481^HeV^) and NiV-G (N481^NiV^) at the primary sequence level. Interestingly, glycosylation at N481^HeV/NiV^ has been shown to modulate fusion activation, suggestive that N502^CedV^ may play a similar role [41]. The otherwise heterogeneous distribution of N-linked glycosylation sequons suggests an absence of functional constraints that would dictate the absolute position of glycan sites across extant HNV-G proteins.

**Figure 2:**
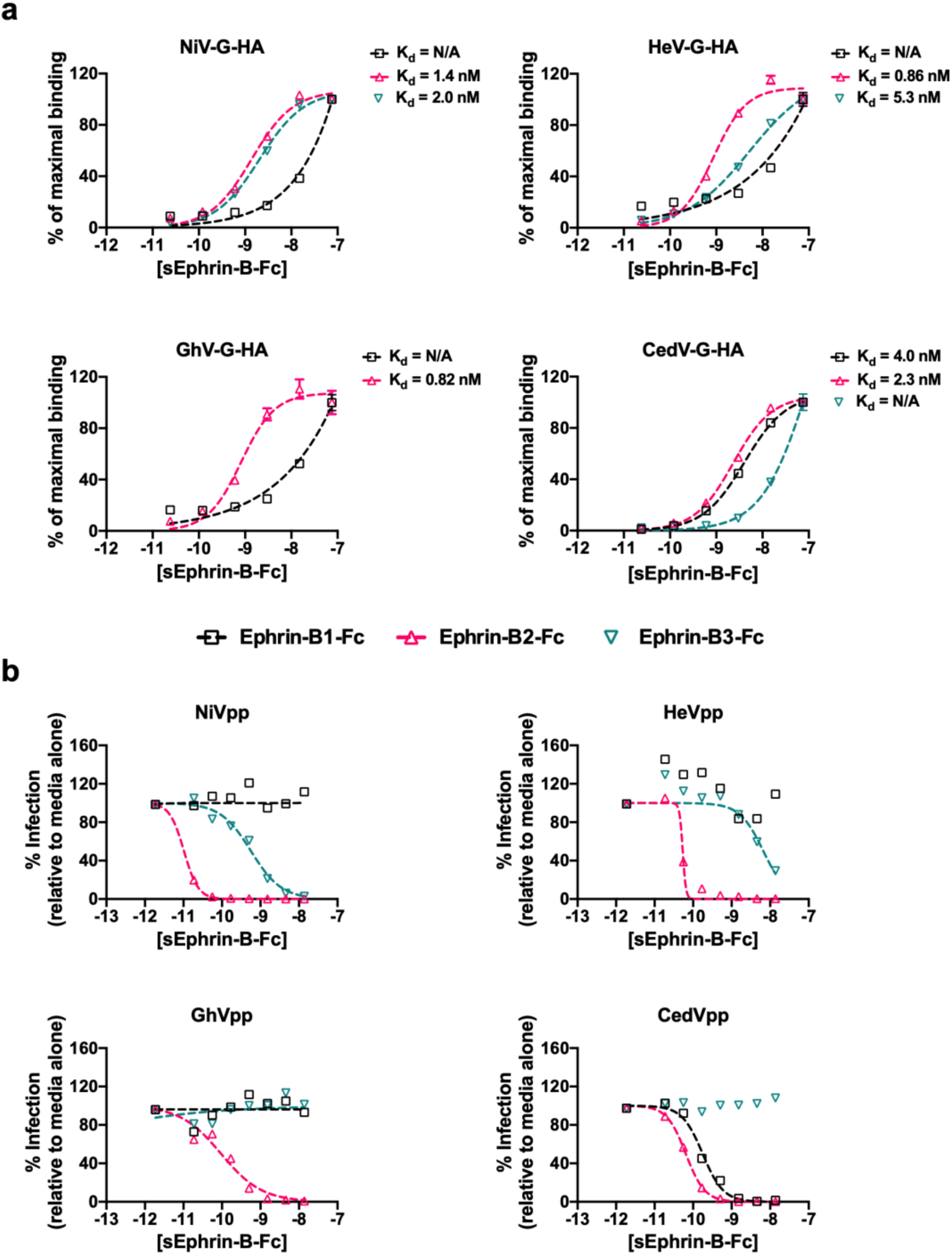
Ephrin-B2 and ephrin-B1 bind CedV-G with nanomolar affinities. **a**, Increasing concentrations of soluble ephrin-B-Fc (10^−11^ to 10^−7^ M) were added to HNV-G transfected HEK-293T cells, and binding (measured as GMFI values) was assessed by flow cytometry using Alexa Fluor 647-labelled anti-human Fc antibodies. Four-parameter dose-response or logistic (4PL) curves were generated by non-linear regression using GraphPad Prism. Data from GMFI values are displayed as percent of maximal binding, with the maximal binding value at the highest concentration of ligand used set to 100%. The bottom of each 4PL curve was constrained to have a constant value of zero. The reported K_d_ (dissociation constant) corresponds to the ephrin-ligand concentration, [sEphrin-B3-Fc], at which 50% maximal binding is achieved. A value of N/A refers to data that could not be fitted unambiguously to a 4PL curve, i.e. binding of soluble ephrin was not titratable or saturation could not be achieved even at concentrations up to 100 nM. Values for ephrin-B3-Fc binding to GhV-G are not displayed, as no detectable binding over background was observed. Data shown are the averages of three independent biological replicates ± SE. **b**, NiV-F/G (NiVpp), HeV-F/G (HeVpp), GhV-F/G (GhVpp), and CedV-F/G (CedVpp) VSV-ΔG-rLuc pseudotyped viruses were used to infect Vero CCL-81 cells in the presence of increasing amounts of soluble ephrin-B1-Fc, ephrin-B2-Fc, and ephrin-B3-Fc fusion proteins. 4PL curves were generated using GraphPad Prism as above. Data from RLU values are displayed as percent of maximal infection, defined as the RLU achieved in the presence of media alone, which is set to 100%. The top and bottom of each 4PL curve was constrained to have a constant value of 100 and 0, respectively. Data shown are the averages of three independent biological replicates ± SE.

Comparison of CedV-G with unliganded HeV-G (Protein Data Bank [PDB] accession code 2X9M) and NiV-G (PDB: 2VWD) reveals considerable structural similarity within the β-propeller scaffold (Cα atom RMSD of 1.7 Å over 385 residues and 1.6Å over 371 residues, respectively) (Fig. 1c), despite substantial primary sequence divergence (Supplementary Fig. 1). Concomitant with genetic proximity, structure-based phylogenetic classification places CedV-G in a cluster of ephrintropic HNV-G proteins, closer to the Asiatic prototype viruses HeV-G and NiV-G than the African GhV-G (Fig. 1d). Such structure-based phylogenetic classification has demonstrable utility in defining clusters of viral receptor-binding glycoproteins that utilize common receptors [18, 42] and, when applied to CedV-G, supports utilization of ephrins. Whilst core secondary structure elements of the β-propeller scaffold are similar, loop regions exhibit substantial structural differences. Interestingly, structural conservation within the apical loops can be broadly divided into two spatially continuous sections: blades β1–3 are structurally variable, and β4–β6 exhibit markedly lower RMSD values (average Cα RMSD of 1.8 Å and 1.0 Å, respectively) (Fig. 1c). Approximately 70% (28/40) of the NiV-G and HeV-G residues that participate in ephrin-B2 recognition are contributed by the structurally conserved β4–β6 blades (Supplementary Fig. 1), suggestive that local structure is constrained by a requirement to maintain ephrin binding.

### CedV-G binds both ephrin-B1 and ephrin-B2 with nanomolar affinity

Given the extent and distribution of structural similarities between CedV-G and ephrin-tropic HNVs, we hypothesized that CedV-G likely utilizes high-affinity ephrin-binding to mediate cellular entry. To this end, we examined the binding of soluble ephrins to cell-surface expressed HNV-G proteins (Fig. 2a). Human embryonic kidney (HEK) 293T cells were transfected with HA-tagged HNV-G glycoproteins (NiV-G, HeV-G, GhV-G, and CedV-G) and titrated against soluble Fc-tagged human B-type ephrins (ephrin-B1-Fc, ephrin-B2-Fc, and ephrin-B3-Fc). In agreement with previous studies [26, 43-45], NiV-G bound both ephrin-B2-Fc and ephrin-B3-Fc with nanomolar affinities (K_d_ =1.4 nM and 2.0 nM, respectively), whilst binding of HeV-G to ephrin-B3-Fc was approximately five-fold weaker than to ephrin-B2-Fc (K_d_ = 5.3 nM and 0.86 nM, respectively) [45, 46]. Unlike the prototypic HNV-Gs, GhV-G exhibited no detectable interaction with ephrin-B3-Fc, but bound strongly to ephrin-B2-Fc (K_d_ = 0.82 nM) [26]. Consistent with our structure-based hypothesis, CedV-G exhibited high-affinity binding to ephrin-B2-Fc (K_d_ = 2.3 nM) but, similar to GhV-G, lacked a titratable interaction with ephrin-B3-Fc. Unexpectedly, we also detected a nanomolar-affinity interaction between CedV-G and ephrin-B1-Fc (K_d_ = 4.0 nM), a receptor with no precedent of HNV-G binding.

As ephrin-B1 binding was unexpected, we sought to determine whether the high-affinity interactions between CedV-G and both ephrin-B1 and ephrin-B2 have relevance to viral entry. We first tested whether cognate soluble ephrin-B ligands could inhibit the entry of HNV pseudotyped particles (HNVpp) into a mammalian cell type that is permissive to HNV infection, Vero-CCL81 cells. HNV envelope glycoproteins (F and G) were pseudotyped onto a recombinant vesicular stomatitis virus (VSV) expressing a *Renilla* Luciferase (rLuc) reporter gene in place of its endogenous envelope glycoproteins (VSV-ΔG-rLuc). Such VSV-based HNVpp, when used as a surrogate in antibody neutralization assays, have been validated by the CDC [47] as equivalent to the gold standard PRNT (plaque reduction neutralization titer) assay using live HNV [11, 26, 43, 44]. We then infected Vero-CCL81 cells with HNVpp in the presence of varying concentrations of soluble ephrin-B-Fc (Fig. 2b). To ensure the veracity of our results, we used a fixed quantity of HNVpp pre-determined to give rLuc activity within the linear dynamic range established for this reporter assay [11]. Entry of both NiVpp and HeVpp was inhibited by soluble ephrin-B2-Fc and ephrin-B3-Fc but not ephrin-B1-Fc (Fig. 2b, top). In contrast, CedVpp entry was inhibited by soluble ephrin-B1-Fc and ephrin-B2-Fc, but not ephrin-B3-Fc, while GhVpp was only inhibited by ephrin-B2-Fc (Fig. 2b, bottom). These entry inhibition data are consistent with the binding data (Fig. 2a) and implicate both ephrin-B1 and ephrin-B2 as receptors for CedV entry.

### Ephrin-B1 and ephrin-B2 support CedV entry

To examine whether ephrin-B1 and ephrin-B2 can function as *bona fide* receptors for CedV entry, we infected ephrin-negative CHO-pgsA745 cells, engineered to stably express ephrin-B1, -B2, or – B3 (CHO-B1, CHO-B2, CHO-B3) [43, 44], with CedVpp or NiVpp. Across several logs of viral input, CedVpp was able to infect both the CHO-B1 and CHO-B2 cells, but not CHO-B3 cells (Fig. 3). Futhermore, NiVpp robustly infected the CHO-B2 and CHO-B3 cells but exhibited markedly reduced entry (1000-fold decrease in RLU) into CHO-B1 cells, as has been previously observed [43]. Thus, ectopic expression of either ephrin-B1 or ephrin-B2 is sufficient to permit entry of CedVpp into an otherwise non-permissive cell-type (CHO-parental) (Fig. 3a-c). Furthermore, CedVpp entry into CHO-B2 cells yielded moderate but consistently higher entry levels than CHO-B1 cells (Fig. 3c versus 3b), corroborating the affinity-binding (Fig. 2a) and ephrin inhibition (Fig. 2b) data, indicating that CedV-G utilizes ephrin-B2 more efficiently than ephrin-B1.

**Figure 3:**
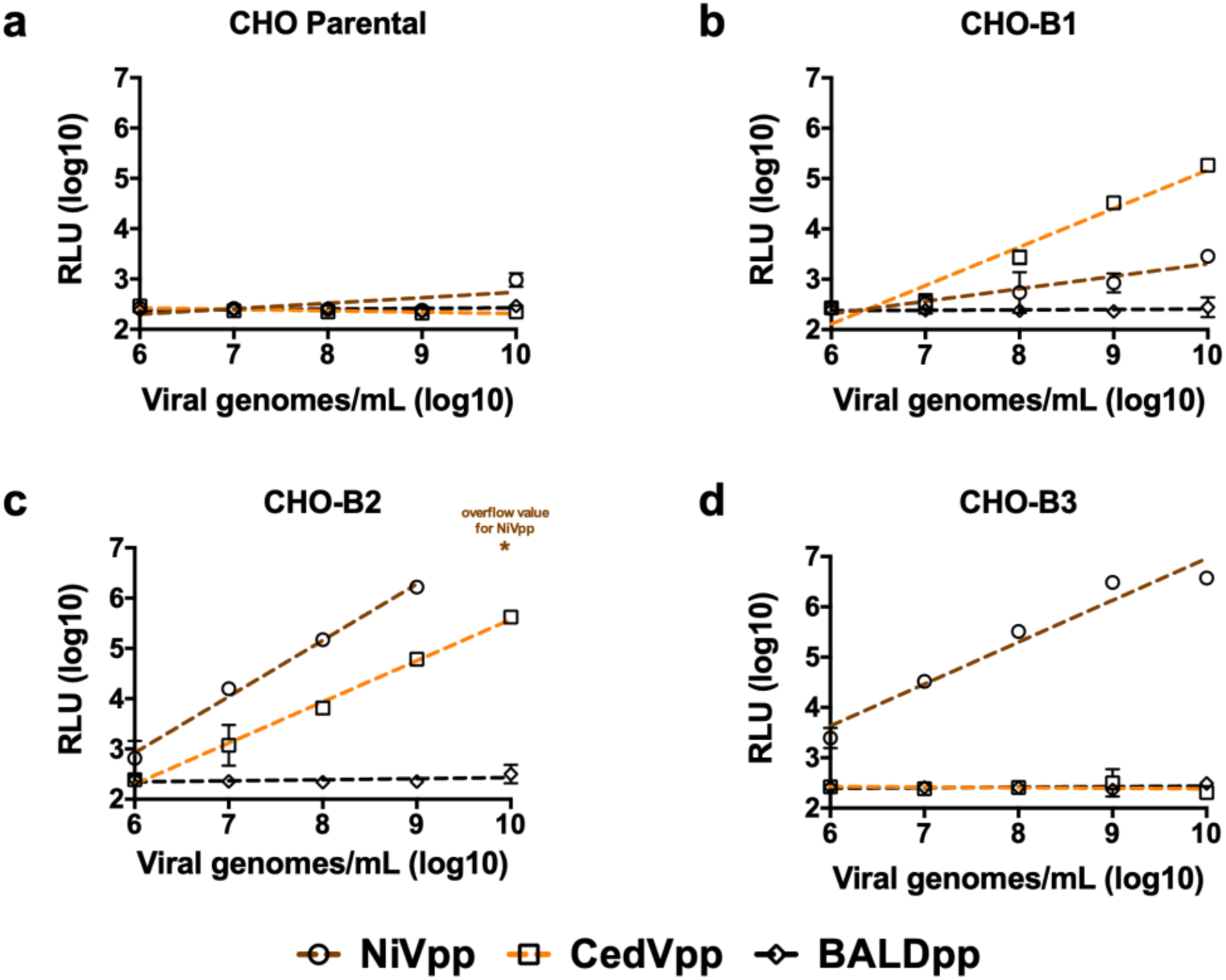
Ectopic expression of ephrin-B1 or ephrin-B2 is sufficient to confer CedVpp entry into a non-susceptible cell type. NiVpp, CedVpp, and BALDpp (VSV pseudotypes bearing no viral glycoprotein) were used to infect **a**, CHO-pgsA745 cells (a naturally ephrin-negative cell line) or CHO-pgsA745 cells that stably express **b-d**, ephrin-B1 (CHO-B1), ephrin-B2 (CHO-B2), or ephrin-B3 (CHO-B3), respectively, over a range of viral inoculum (viral genomes/mL). Entry was measured as described in the legend to Figure 2. The asterisk indicates an RLU value above the maximum limit of detection. RLU were plotted against numbers of viral genome copies per milliliter and fitted to a linear regression (dashed lines) using GraphPad Prism. Data shown are the averages of three independent biological replicates ± SE.

Intriguingly, although both NiV-G and CedV-G exhibit nanomolar affinity binding to ephrin-B2 (Fig. 2a), approximately an order of magnitude more CedVpp viral genomes (per mL) were required to achieve entry levels equivalent to NiVpp on CHO-B2 cells (Fig. 3c). To eliminate the possibility that NiV glycoproteins are better incorporated into VSV pseudotypes than the CedV equivalents, we performed a western blot analysis, which demonstrated that neither CedV-G nor the fusion-competent cleavage product of CedV-F (CedV-F_1_) were significantly less incorporated (Supplementary Fig. 3a). Thus, the reduced infectivity of CedVpp is likely a consequence of intrinsically reduced fusogenicity of CedV-F/G compared to NiV-F/G, as evidenced by smaller and fewer syncytia formed by CedV-F/G relative to both NiV-F/G and HeV-F/G, in highly permissive U87 glioblastoma cells (Supplementary Fig. 3b). Despite the lower fusogenicity of CedV-F/G and reduced infectivity, relative to NiVpp on CHO-B2 cells, CedVpp infection was consistently 1–2 logs higher than NiVpp on CHO-B1 cells (Fig. 3b). Only at the highest level of viral inoculums (>10^9^ viral genomes/ml) did NiVpp exhibit very low levels of infectivity on CHO-B1 cells. Altogether, these data indicate that CedV is unique amongst known HNVs in its ability to specifically utilize both ephrin-B1 and ephrin-B2 for cellular entry.

### Ephrin-B1 supports CedV entry into physiologically relevant primary cells

As endothelial cells constitute the primary targets of HNV infection *in vivo* [48-50], we utilized primary human umbilical vein endothelial cells (HUVECs) to investigate the importance of ephrin-B1 as a functional entry receptor for CedV in a physiologically relevant cell type. Real-time quantitative PCR (qPCR) analysis of B-class ephrin transcripts within primary HUVECs revealed the presence of mRNA encoding both ephrin-B1 and ephrin-B2, whereas ephrin-B3 mRNA was undetectable (Fig. 4a). Following confirmation of active transcription of ephrin-B1 and ephrin-B2 in HUVECs, soluble envelope competition assays were performed to assess HNVpp entry (Fig. 4b-c).

**Figure 4:**
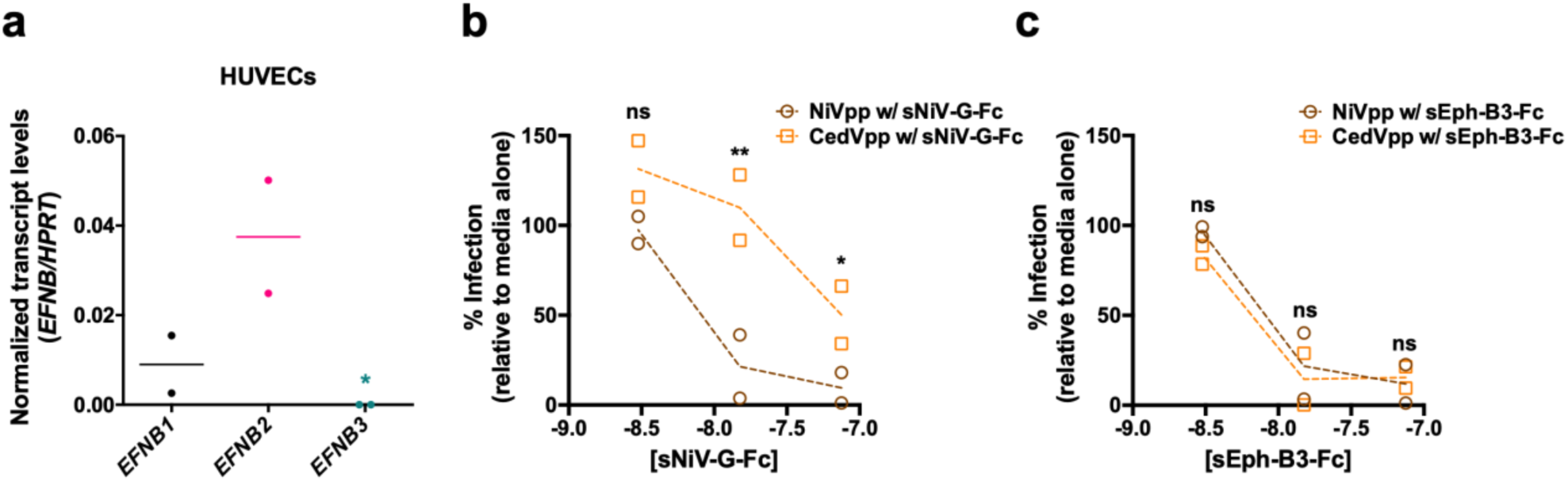
Ephrin-B1 facilitates CedVpp entry into biologically relevant primary HUVECs. **a**, Active transcription of ephrin-B1, ephrin-B2, and ephrin-B3 in primary human umbilical vein endothelial cells (HUVECs) was determined by quantitative PCR. Transcript levels are shown normalized to hypoxanthine phosphoribosyltransferase (HPRT) transcripts. Ephrin-B3 transcript levels were below the limit of detection (asterisk). Data shown are the individual data points from two independent biological replicates. Horizontal dashes represent the mean from the two replicates. **b**, NiVpp, CedVpp, and VSVpp were used to infect primary HUVECs in the presence of increasing amounts of soluble Fc-tagged NiV-G (sNiV-G-Fc) or soluble Eph-B3 receptor (sEph-B3-Fc). Entry was measured as in Figure 2. Data are shown as percent infection relative to the signal achieved when the viruses are incubated in the presence of media alone. Data shown are the individual data points from two independent biological replicates. Dashed lines connect the means from the duplicate data. Statistical significance for this entry inhibition assay was tested with a two-way ANOVA with Holm-Sidak’s correction for multiple comparisons, n/s denotes no significance, * denotes p<0.05, ** denotes p<0.005.

Pre-incubation of HUVECs with saturating quantities of sNiV-G-Fc completely inhibited NiVpp entry, suggesting that all available cell-surface displayed ephrin-B2 molecules were sequestered by soluble NiV-G and thus viral entry was not supported in the absence of an alternate cognate receptor for NiV, namely ephrin-B3. Conversely, under identical conditions, CedVpp entry was still supported at ∼50%, evidencing ephrin-B1-mediated entry of CedVpp in the absence of available ephrin-B2 (Fig. 4b). Entry of both NiVpp and CedVpp was completely abrogated by pre-incubation with soluble Eph-B3 receptor (sEph-B3-Fc), an Eph receptor that binds to all three B-class ephrin ligands with similar affinities [51], further confirming that viral entry is ephrin-dependent (Fig. 4c). These data reveal that endogenously-expressed ephrin-B1 supports CedVpp entry in a physiologically relevant cell type, validating the role of ephrin-B1 as a functional entry receptor for CedV.

### The CedV-G–ephrin-B1 structure reveals a conserved HNV-G–ephrin interaction mode

Following our findings that CedV was able to utilize a previously unreported receptor repertoire, we sought to delineate the molecular features of ephrin binding and ligand selectivity. To this end, we solved the crystal structure of the attachment-mediating β-propeller of CedV-G in complex with the extracellular β-barrel domain of human ephrin-B1, to 4.07-Å resolution. Five complexes populated the crystallographic asymmetric unit and displayed no appreciable structural variation, within the limits of the resolution. Analyses herein concern the complex comprising chains ‘B’ and ‘D’ which was selected for superior quality electron density and the model quality permitted as a consequence.

Comparison with the unliganded structure of CedV-G reveals little overall structural variation (0.4 Å RMSD over 417 equivalent Cα) within the ephrin-B1-bound CedV-G scaffold. Notably, the β6-S2–S3 loop, which is conformationally distinct in each of the unliganded CedV-G crystallographic asymmetric unit (a.s.u.) copies (Fig. 1a), is also the region of greatest structural variability when comparing the ephrin-B1 bound and unbound states of CedV-G. Plasticity and ephrin ligand-induced structural transitions within this region mirror observations in other HNV-G proteins and their ephrin-bound complexes [36, 37].

CedV-G engages ephrin-B1 with an overall binding mode that is similar to that utilized by other HNV-G proteins, when recognizing their respective ephrin receptors (Fig. 5a-c)[26, 45, 46]. CedV-G and ephrin-B1 form a 1:1 complex with an extensive molecular interface (buried surface area: 2,900 Å^2^), which is larger than previously characterized HNV-G–ephrin interfaces (NiV-G– ephrin-B2 = 2,800 Å^2^; NiV-G–ephrin-B3 = 2,700 Å^2^; HeV-G–ephrin-B2 = 2,600 Å^2^; GhV-G–ephrin-B2 = 2,300 Å^2^, calculated using the PDBePISA server [52]). Ephrin-B1 is bound at the top of the molecule and forms an interface that is chiefly comprised of residues from the structurally conserved β4–6 blades of CedV-G, which contribute ∼70% (∼2,000 Å^2^) of the total binding surface area (Figs. 1, 5c, and Supplementary Fig. 1). Despite localizing to the interface, the glycan at N425 extends away from ephrin-B1 and does not appear to participate in receptor binding.

**Figure 5:**
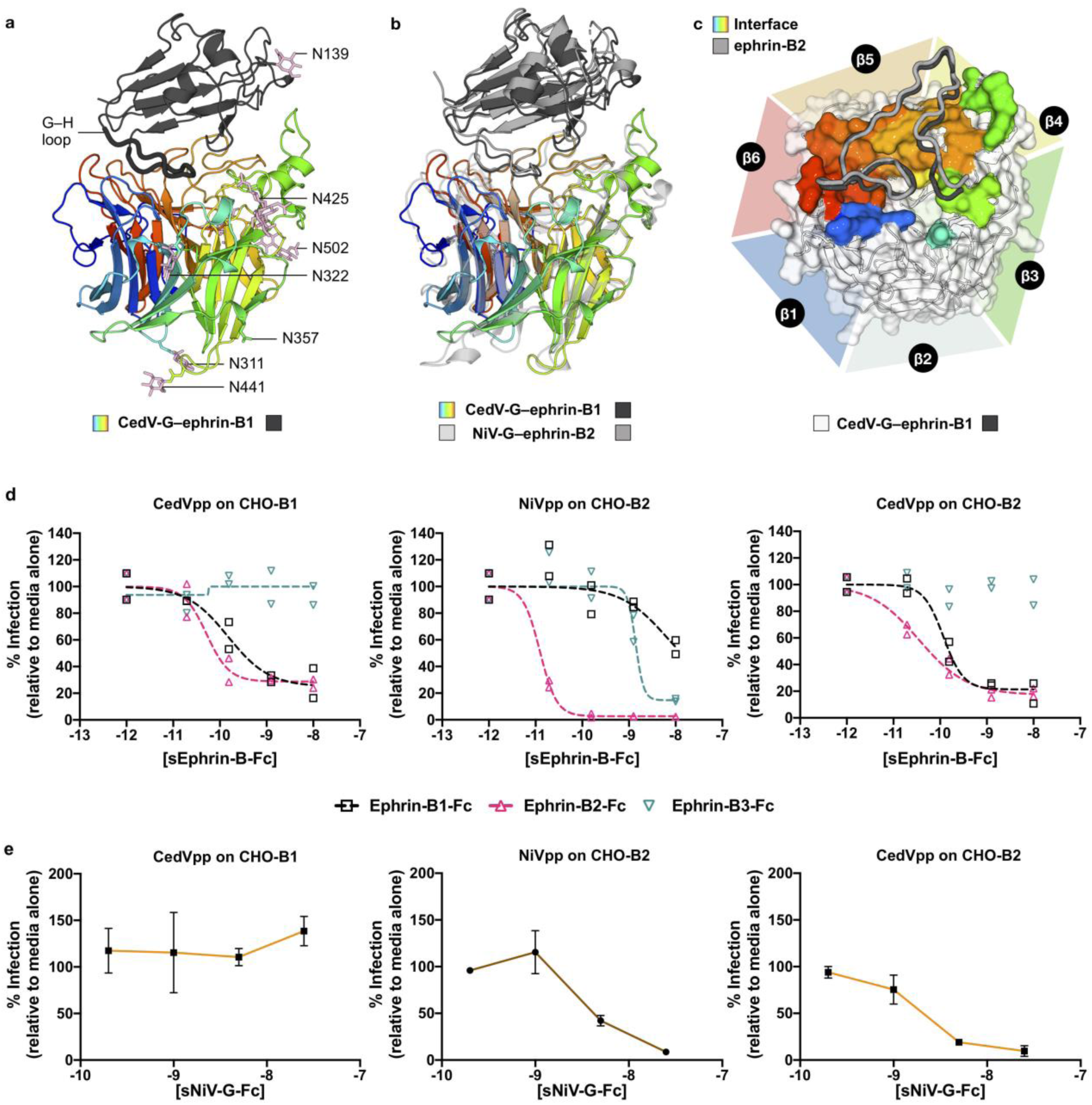
CedV-G binds ephrin-B1 and ephrin-B2 at a conserved and overlapping binding site. **a**, The crystal structure of CedV-G in complex with ephrin-B1 reveals a conserved mode of receptor engagement across ephrin-tropic HNV-G proteins. The receptor-binding domain of CedV-G (colored in a gradient from blue to red, from N- to C-terminus) forms a 1:1 complex with the extracellular domain of ephrin-B1 (dark grey). The principal interaction region of ephrin-B1, the ‘G–H loop’, is displayed as a thick tube for clarity. Modelled N-linked glycan moieties (pink) and the asparagine residues of all putative N-linked glycosylation sequons are shown as sticks. **b**, Superposition of NiV-G–ephrin-B2 (PDB: 2VSM) on CedV-G–ephrin-B1. The NiV-G–ephrin-B2 complex is colored in light grey, with NiV-G shown as transparent for clarity. **c**, Comparison of bound ephrin-B1 and ephrin-B2 molecules. CedV-G is shown as a white transparent surface with the ephrin-B1 interface colored according to sequence position as in panel **a.** Regions of ephrin-B1 (dark grey) and ephrin-B2 (light grey) bound by CedV-G and NiV-G (respectively) are shown as a cartoon tubes. β-propeller blades are delineated with triangles that are colored according to sequence position as in panel **a. d**, Ephrin-B1 and ephrin-B2 compete for binding to CedV-G. NiVpp and/or CedVpp were used to infect CHO-B2 cells (*middle and right panels*) or CHO-B1 cells (*left panel*), as indicated, in the presence of increasing amounts of soluble ephrin-B1-, ephrin-B2-, and ephrin-B3-Fc. Entry was measured as in Figure 2. 4PL dose-response curves were generated as in Figure 2 and based on values displayed as percentages of infection with the RLU achieved in the presence of media alone set to 100%. The top of each fit curve was constrained to a have a constant value of 100. Data shown are the individual data points from two independent biological replicates with the fit curves shown in dashed lines. **e**, NiV-G and CedV-G compete for binding to ephrin-B2. NiVpp or CedVpp were used to infect CHO-B2 *(middle and right panels)* or CHO-B1 *(left panel)* cells in the presence of increasing amounts of soluble Fc-tagged NiV-G (sNiV-G-Fc). Data are shown as percent infection relative to the signal achieved when the viruses were incubated in the presence of media alone. Data shown are the averages of three independent biological replicates ± SE.

Like the other HNV-G–ephrin complexes, the G–H loop of ephrin-B1 (residues 113–129^ephrin-B1^) constitutes a major interacting region and is inserted into the central depression on top of CedV-G, contributing 1,700-Å^2^ BSA to the molecular interface (Fig. 5a–c). Interestingly, despite sequence variation and pronounced structural dissimilarity in their unliganded states [53], the G–H loops of ephrin-B1 and ephrin-B2 adopt a strikingly similar conformation in their HNV-G-bound states (Fig. 5c). Indeed, the Greek-key fold of the bound ephrin-B1 ectodomain is highly similar to that of NiV-G-bound ephrin-B2 (1.0-Å RMSD across 135 aligned Cα atoms), though less similar to ephrin-B3 (1.6 Å RMSD across 126 aligned Cα atoms). Together, binding-induced structural transitions within both CedV-G and the ephrin G–H loops support a model of an induced-fit mechanism of ephrin recognition that is conserved across ephrin-tropic HNVs[36, 37, 45].

### CedV-G utilizes a conserved mode of ephrin recognition

Given the striking structural similarities between CedV-G–ephrin-B1 and other HNV-G-ephrin complexes (Fig. 5b), we hypothesized that CedV-G likely utilizes the same receptor binding site to engage ephrin-B2. To assess this, we determined pseudotyped virus entry into CHO-B1 and CHO-B2 cells in the presence of competing soluble B-class ephrin ligands. As expected[43, 54, 55], ephrin-B2-Fc and ephrin-B3-Fc inhibited NiVpp entry into CHO-B2 cells, while ephrin-B1-Fc failed to strongly inhibit entry at concentrations as high as 10 nM (Fig. 5d, middle panel), confirming that ephrin-B2 and ephrin-B3 are each bound by the same site on NiV-G [45]. Similarly, both ephrin-B2-Fc and ephrin-B1-Fc inhibited CedVpp entry into CHO-B2 cells (Fig. 5d, right panel), evidencing the ability of ephrin-B1 to block ephrin-B2-dependent CedVpp entry through competition for an overlapping binding site on CedV-G. Moreover, ephrin-B2-Fc inhibited CedVpp entry into CHO-B1 cells (Fig. 5d, left panel). In both CHO-B2 and CHO-B1 cells, ephrin-B2-Fc-mediated inhibition of CedV-G was more potent than ephrin-B1-Fc (Fig. 5d), further supporting our binding (Fig. 2) and entry (Fig. 3) data that suggest ephrin-B2 is more efficiently utilized than ephrin-B1.

To further validate our hypothesis, we performed soluble envelope competition assays in which CHO-B2 cells were incubated with soluble NiV-G-Fc, prior to infection with NiVpp or CedVpp (Fig. 5e, middle and right panels, respectively). Entry of NiVpp and CedVpp on CHO-B2 cells was completely inhibited by NiV-G-Fc, indicative that CedV-G and NiV-G recognize a common interface on ephrin-B2, which, in conjunction with the structures of CedV-G (Fig. 1) and CedV-G– ephrin-B1 (Fig. 5a–c), support a universal mode of ephrin recognition across ephrin-tropic HNVs. Importantly, CedVpp entry into CHO-B1 cells was unaffected by saturating amounts of NiV-G-Fc (Fig. 5e, left panel). Thus, despite a shared mode of receptor engagement across ephrin-tropic HNVs, these data indicate that CedV-G possesses subtle but distinct features within its receptor binding site that determine utilization of its idiosyncratic receptor repertoire.

### Accommodation of the YM motif of ephrin-B1 is a key determinant of receptor specificity

In order to determine the molecular features that underscore the distinct ephrin specificity of CedV-G, we examined both the structural and functional implications of two hydrophobic motifs that differ between ephrin-B1 and ephrin-B2. Whilst the G–H loops of B-type ephrin paralogs are highly conserved (Fig. 6a), ephrin-B1 and ephrin-B2 differ by the presence of a Tyr^121^–Met^122^ (YM) or Leu^121^–Trp^122^ (LW) motif, which have previously been shown to be critical determinants of differential receptor utilization [43]. Inspection of the CedV-G–ephrin-B1 complex structure reveals that the side chain of Tyr^121-ephrin-B1^ is inserted into a large hydrophobic cavity that is unique to CedV-G (Fig. 6b-c). Indeed, the equivalent region within NiV-G (Trp^504^), HeV-G (Trp^504^), and GhV-G (Trp^514^) is, in all instances, occluded by a tryptophan side chain that protrudes toward the toroidal axis of the β-propeller, rather than being sequestered in the opposing direction as is the equivalent residue, Tyr^525-CedV-G^ (Fig. 6c). Thus, it is likely that the expanded hydrophobic cavity of CedV-G is critical to ephrin-B1 recognition as it circumvents steric clashes that would preclude its recognition by other HNV-Gs.

**Figure 6:**
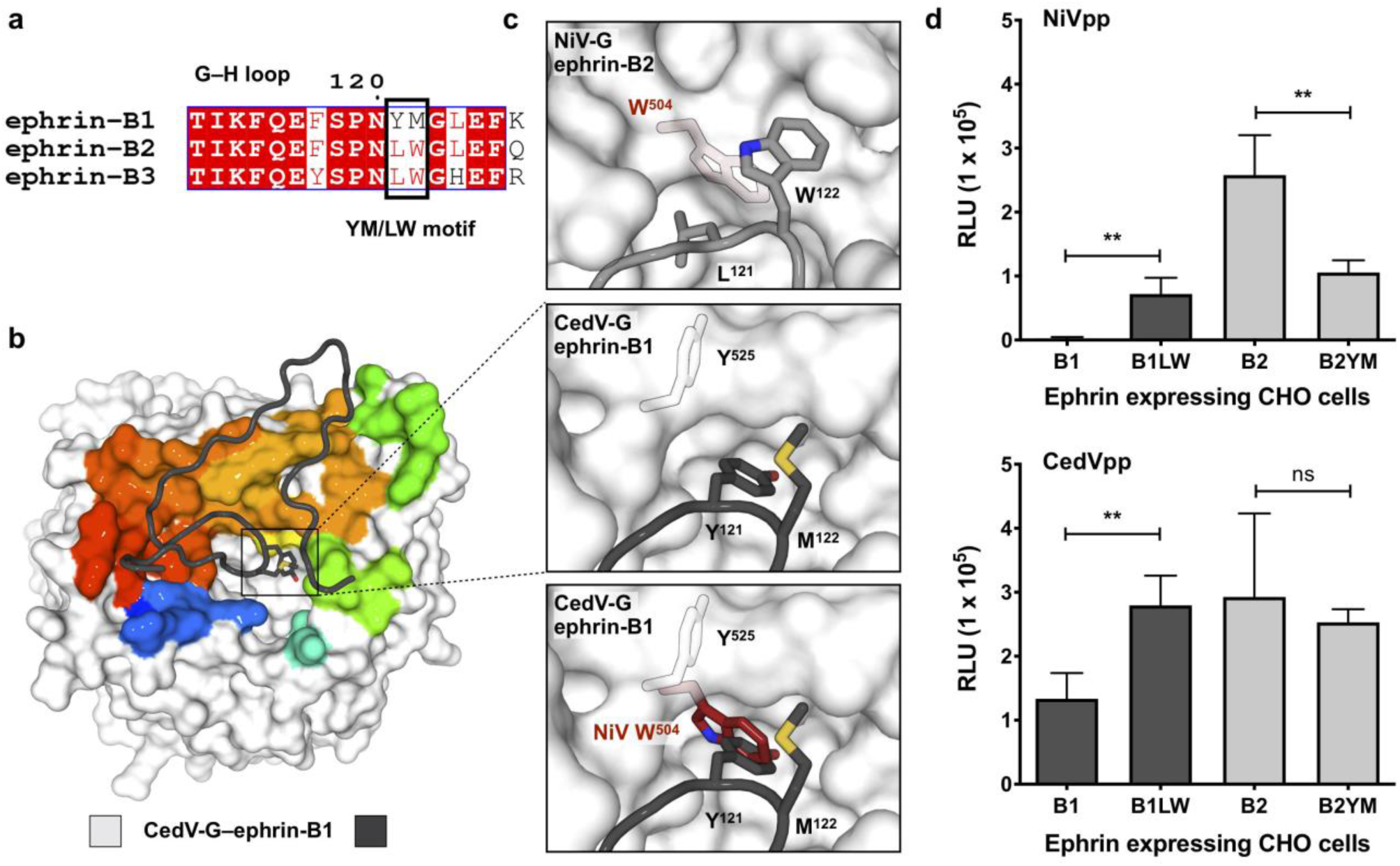
Accommodating the YM motif is critical for ephrin-B1 utilization by CedV. **a**, Sequence alignment of the G–H loop region of human B-type ephrins. Absolutely conserved residues are highlighted red, partially conserved residues are colored red, and non-conserved residues are black. Residues that constitute the YM/LW motif are outlined with a black box. Sequences are numbered according to ephrin-B1. Alignments were determined by MultAlin [78] and plotted using ESPript [79]. **b**, Structure of CedV-G–ephrin-B1. CedV-G is displayed as a white surface with the ephrin-B1 interface colored according to sequence position (blue to red, N- to C-terminus). The principal interacting region of ephrin-B1 is shown as a dark grey cartoon tube, with the side chains of Y121 and M122, the ‘YM motif’, shown as sticks and colored according to constituent atoms. **c**, Detailed view of the YM motif of ephrin-B1 (boxed region in **a**; *middle*) and the equivalent view of the LW motif of NiV-G-bound ephrin-B2 (light grey; *top*). (*bottom*) Overlay of the NiV-G residue, W504 (red) onto CedV-G–ephrin-B1, demonstrates potential steric overlap between NiV-G and ephrin-B1. The side chain of Y525 in CedV-G is sequestered away from the receptor binding site. Side chains of key residues are shown as sticks and colored according to constituent atoms. **d**, CedV is more tolerant to substitution of the YM/LW motif than NiV. CHO cells expressing both wild-type ephrins (B1 and B2) and mutants with reciprocally exchanged LW/YM motifs (B1LW and B2YM) were infected with NiVpp (*top*) or CedVpp (*bottom*). Entry was assessed and quantified as in Fig. 2. Data represent the average of quadruplicate measurements ± SE. Statistical significance for the indicated comparisons were evaluated using a two-tailed unpaired t-test, ** denotes p<0.005 and n/s denotes no significance.

To interrogate this structure-based hypothesis, we tested pseudotyped viral entry into CHO-pgsA745 cells stably expressing a panel of wild-type and mutant ephrin ligands (Fig. 6d). These stable cell lines had been previously demonstrated to express similar levels of the indicated wild-type and mutant ephrin ligands, as measured by sEph-B3-Fc binding [43]. As anticipated, NiVpp and CedVpp were able to enter cells expressing ephrin-B2_WT_, whilst ephrin-B1_WT_ supported only CedVpp entry. Replacement of the ephrin-B2 LW motif with the equivalent YM residues of ephrin-B1 (ephrin-B2_YM_) severely impaired the ability of ephrin-B2_YM_ to support NiVpp entry, though did not prevent it entirely. Conversely, the reciprocal exchange of the YM to LW motif in ephrin-B1 (ephrin-B1LW) rendered it capable of supporting some level of NiV entry. Intriguingly, ephrin-B1_LW_-mediated entry of CedVpp was increased relative to ephrin-B1_WT_, suggestive that the YM motif is less efficiently utilized than LW for CedV entry, in the context of an ephrin-B1 background. By contrast, introduction of the YM motif into ephrin-B2 induced no appreciable difference in its ability to support CedVpp entry. The differential response to the YM/LW motifs, when presented in the context of distinct ephrin backgrounds, highlights the importance of the myriad surrounding interactions that comprise the extensive protein–protein interface. Taken together, this integrated structural and functional analysis suggests that the ability of CedV-G to effectively accommodate the YM motif of ephrin-B1, in a sterically unrestrained cavity, is a critical determinant that directs ephrin-B1 utilization.

## Discussion

Viral surveillance continues to revise the known geographic coverage and biological diversity of HNV species [12]. Whilst identification of HeV and NiV was a consequence of their emergence as the etiological agents of severe respiratory and neurological disease, targeted virus discovery efforts have demonstrated utility in identifying novel HNVs yet to be linked with symptomatic illness[10, 11, 19]. One such virus, CedV, represents only the third HNV species to be isolated, and remains the sole member of the genera confirmed as non-pathogenic [18, 19]. Combined, the starkly contrasting virulence and the ecological and biological similarities between CedV and the deadly HNVs demarcates CedV as a valuable model with which to assess HNV functional diversity and the determinants of pathogenesis.

Through adoption of a combined structural and functional approach, we sought to determine the receptor repertoire utilized by CedV-G and uncover molecular features that dictate its specificity. We demonstrate that CedV-G utilizes a structurally constrained receptor-binding architecture to mediate recognition of highly conserved ephrin receptors utilizing an analysis that relates local structural conservation to discrete regions that are functionally constrained amongst genetically diverse HNVs (Fig. 1c). By demonstrating ephrin-B2 mediated cellular entry of CedV, we confirm previous studies [19, 35] and provide evidence for a conserved receptor recognition mode (Fig. 5). Unexpectedly, we detected high-affinity binding to ephrin-B1, a B-class ephrin with no precedent of supporting HNV entry (Figs. 2 and 3) and demonstrate that the interaction is sufficient to permit viral entry into pathobiologically relevant primary endothelial cells (Fig. 4). Furthermore, we present crystal structures of CedV-G, in both unbound and ephrin-B1-bound states (Figs. 1 and 5), which reveal a universal mode of receptor recognition across distinct B-type ephrin ligands and diverse HNV-G proteins.

The bipartite nature of structural conservation within CedV-G (Fig. 1c) is indicative of contrasting selective pressures which, in blades β1–β3 permit diversification, but in the β4–β6 blades, constrain variation to maintain high-affinity ephrin binding. Like NiV and HeV, CedV utilizes ephrin-B2 more efficiently than its cognate alternate receptor, ephrin-B1 (or ephrin-B3 for NiV and HeV) (Figs. 2 and 3) [43, 45]. Furthermore, despite its idiosyncratic receptor repertoire, structure-based phylogenetic analysis co-localizes CedV-G with other ephrin-tropic HNV-Gs, all of which are unified in utilizing ephrin-B2 for cellular entry (Fig. 1d). Combined, the more effective ephrin-B2 usage and structure-based phylogenetic association of all ephrin-tropic HNV-Gs implicate ephrin-B2 as a receptor utilized by ancestral HNVs, alongside which coincident alternate receptor specificities may have arisen.

Despite evident commonalities of HNV-G-mediated ephrin recognition, the structure of CedV-G–ephrin-B1 reveals unique molecular properties that provide a structure-based rationale for receptor specificity. Modulation of the side chain orientation of a single amino acid in CedV-G (Y^525^) relieves a steric barrier that likely precludes ephrin-B1 recognition by other HNV-Gs. Given that only minor changes to ephrin-B1 are required for it to support NiV entry (Fig. 6d) [43], and the existence of relatively subtle structural differences within HNV-G receptor binding sites, it is possible that the existence or acquisition of ephrin-B1 tropism in extant HNVs may not be uncommon. Although CedV-G is unable to utilize ephrin-B3, our structural hypothesis suggests that acquired ephrin-B1 specificity does not necessarily come at the expense of ephrin-B3 usage, as the LW motif is common to both ephrin-B2 and ephrin-B3.

The acquisition of alternate receptor specificities, and the species and cell-type tropism thereby engendered, is of acute biomedical and agricultural importance [27]. For example, the systemic dissemination and multi-organ vasculitis associated with NiV and HeV is consistent with expression of ephrin-B2 in endothelial cells [54, 56] and peripheral organs, such as the kidney, lung, and spleen, wherein ephrin-B3 is less abundant. However, central nervous system (CNS) disease pathologies [57, 58], which are the ultimate cause of death in fatal NiV and HeV infection, are likely a consequence of markedly increased and specific expression of ephrin-B3 in multiple brain regions (Supplementary Fig. 5) [59-61].

Although the absence of interferon-antagonism in CedV likely plays a critical role in determining its apathogenic phenotype [19, 20], an inability to utilize ephrin-B3 may also be a contributing factor. Interestingly, although ephrin-B1 is expressed at low to negligible levels in the CNS, its expression in other tissues is both more widespread and often greater in magnitude than ephrin-B2. Of note is the relatively high expression of ephrin-B1 in the lung, esophagus, and salivary glands (Supplementary Fig. 5), which suggests that ephrin-B1 utilization could augment aspects of oropharyngeal transmission postulated for HNVs [62]. Thus, whilst the pathobiological and ecological implications of ephrin-B1 tropism are presently unclear, our study sets a precedent for ephrin-B1 utilization, and in so doing expands the known repertoire of HNV cellular entry receptors utilized by this group of lethal human pathogens. Finally, in light of our structure-function analyses, it is plausible that the barrier for ephrin-B1 usage may not be high, and that other HNVs may acquire, or have already acquired, an expanded repertoire of B-class ephrin receptors that could modulate pathogenicity and transmissibility. Characterization of receptor tropism characteristics should be central to future surveillance efforts that aim to identify and assess new HNVs, in order to elucidate the association between pathogenicity and expanded receptor usage.

## Materials and Methods

### Protein production

Independent expression vectors for the putative six-bladed β-propeller domain of CedV-G (residues 209–622; NCBI reference sequence: YP_009094086.1) and the G-interacting N-terminal extracellular domain of ephrin-B1 (residues 29–167; NCBI reference sequence NP_004420.1) were generated by PCR-amplification from codon-optimized synthetic cDNA templates (GeneArt, Life Technologies) and subsequent restriction-based cloning into the pHLsec mammalian expression vector [63].

HEK-293T cells (ATCC CRL-1573) were transiently transfected with the desired protein constructs and expressed in the presence of the class 1 α-mannosidase inhibitor, kifunensine [64]. For the generation of CedV-G–ephrin-B1 complexes, cells were co-transfected with a 1:1.5 mass ratio of CedV-G to ephrin-B1 cDNA. Cell supernatants were harvested and clarified 72 h post transfection, after which supernatants were diafiltrated against a buffer containing 10 mM Tris (pH 8.0) and 150 mM NaCl (ÄKTA Flux diafiltration system; GE Healthcare). In all instances, glycoproteins and their complexes were purified by a tandem immobilized nickel affinity and size exclusion chromatography (SEC) strategy utilizing HisTrap™ HP and Superdex™ increase 200 10/300 GL columns (GE Healthcare), equilibrated in 10 mM Tris (pH 8.0) and 150 mM NaCl, respectively. To aid crystallogenesis of unliganded CedV-G high-mannose-type N-linked glycans were trimmed by partial enzymatic deglycosylation with endoglycosidase F1 (25°C for 18 h).

For the production of Fc-tagged HNV-G proteins utilized in cellular assays, codon-optimized DNA fragments corresponding to the receptor-binding domains of NiV-G (residues 183–602; NCBI reference sequence: NC_002728.1) and CedV-G (residues 209–622; NCBI reference sequence: YP_009094086.1) were cloned into the pHL-FcHis vector [63]. Transfection and protein purification procedures were identical to those utilized for the production of non-Fc-tagged proteins, detailed above. In all instances, kifunensine was not presence during the expression of HNV-G-Fc.

### Crystallization and structure determination

Endoglycosidase F1-treated CedV-G was concentrated to 6.7 mg/ml and subjected room temperature crystallization screening utilizing the sitting-drop vapor diffusion method [65]. A single crystal of CedV-G was observed after 15 days in a condition comprising 5% (w/v) polyethylene glycol 6000 and 0.1 M MES pH 6.0. Additionally, CedV-G–ephrin-B1 containing high-mannose glycans derived from expression in the presence of kifunensine, was concentrated to 12.2 mg/ml. Crystals formed after 132 days in a condition comprising 25% polyethylene glycol 1500 (w/v), 0.1 M PCB (sodium propionate, sodium cacodylate, and BIS-TRIS propane) pH 5.0 and 0.02 M cadmium chloride dihydrate [66].

Crystals were harvested and cryo-protected by transfer into a drop of the respective mother liquor supplemented with 25% glycerol (v/v) prior to flash cooling in liquid nitrogen. X-ray diffraction data were collected for CedV-G (to 2.78-Å resolution) and CedV-G–ephrin-B1 (to 4.07-Å resolution) at beamlines I02 and I24 at Diamond Light Source (UK), respectively. Crystal data were indexed, integrated, and scaled using XIA2 [67]. The structure of CedV-G was solved by molecular replacement, implemented in PHASER [68], using the structure of unliganded NiV-G (PDB accession code 2VWD) as a search model. The refined structure of CedV-G and ephrin-B2, derived from the NiV-G-ephrin-B2 complex (PDB: 2VSM), were used as independent search models to solve the structure of CedV-G–ephrin-B1. In both instances, iterative rounds of model building and refinement were performed using COOT [69], PHENIX [70], and REFMAC [71], utilizing non-crystallographic symmetry restraints and TLS parameterization. The structure of CedV-G–ephrin-B1 was refined using grouped B-factors and reference model restraints derived from the higher resolution CedV-G structure. Conformational validation of N-linked glycans was performed using Privateer [72]. Data collection and refinement statistics are presented in Supplementary Table 1.

### Structure-based phylogenetic analysis

Available structures of unique paramyxovirus receptor-binding glycoproteins (G/H/HN) were utilized to construct a structure-based phylogenetic tree. Structures used were as follows: Human Parainfluenza virus 3, HPIV3-HN (PDB accession code: 1V3B) [73]; Newcastle disease virus, NDV-HN (1E8T) [74]; Parainfluenza virus 5, PIV5-HN (4JF7) [40]; Mumps virus, MuV-HN (5B2C) [75]; Ghana virus, GhV-G (4UF7) [26]; Nipah virus, NiV-G (2VWD) [36]; Hendra virus, HeV-G (2X9M) [37]; Cedar virus, CedV-G; Measles virus, MeV-H (2RKC); Mòjiāng virus, MojV-G (5NOP) [18]. A pairwise evolutionary distance matrix was calculated using the Structural Homology Program (SHP) [76] and plotted as an unrooted phylogenetic tree using PHYLIP [77].

### Cells and culture conditions

HEK-293T, Vero-CCL81, and U87 (ATCC) cells were grown in DMEM supplemented with 10% FBS and 1% penicillin/streptomycin (Thermo). Previously described CHO-pgsA745 cells expressing either wild-type or mutant ephrins [43] were maintained in DMEM/F12 medium (Thermo) with 10% FBS and 1% penicillin/streptomycin. Human umbilical vein endothelial cells (pooled donors; Lonza) were maintained in EndoGRO-LS Complete Media Kit, composed of EndoGRO Basal Medium and the EndoGRO Supplement Kit (Millipore Sigma). EndGRO media was also supplemented with 2% FBS, 1% penicillin/streptomycin, and 1% GlutaMAX supplement (Thermo). HUVECs utilized for soluble envelope competition experiments were between passage three and six.

### Plasmids and reagents

Codon-optimized sequences of CedV-G (GenBank accession number AJP33320.1) and CedV-F (GenBank accession number YP_009094085.1) were tagged with a C-terminal HA or AU1, respectively, and cloned into pCAGGS mammalian expression vectors, as previously described for extant HNV glycoproteins [26, 44]. Soluble human ephrin-B1-Fc, ephrin-B2-Fc, ephrin-B3-Fc, and Eph-B3-Fc utilized for competition experiments were purchased from R&D Systems.

### Cell surface binding assay

Binding of soluble Fc-tagged human ephrin-B ligands to HNV-G transfected cells was assessed by flow cytometry. HEK-293T cells were transfected, using Lipofectamine 2000 (Thermo), with equal concentrations of plasmids encoding the indicated HNV-G proteins, or an empty vector control. Transfected cells were subsequently double stained with a primary rabbit anti-HA polyclonal antibody (pAb) (Novus, catalogue number: NB600-363) at a 1:1,000 dilution, as well as increasing amounts of the ephrin-B1-Fc, ephrin-B2-Fc, and ephrin-B3-Fc ligands for 1 h at 4 ^°C^. Cells were then washed with buffer (2% FBS/PBS) and incubated with an Alexa 488-labelled goat anti-rabbit antibody (Thermo, catalogue number: A-11034) diluted 1:2,000, to identify HNV-G positive cells, and an Alexa 647-labelled goat anti-human antibody (Thermo, catalogue number: A-21445) diluted 1:2,000, to capture ephrin-B binding, for 1 h at 4 °C. Cells were washed in 2% FBS/PBS, fixed in 2% PFA/PBS, and subjected to flow cytometry (Guava easyCyte).

Data were analyzed using FlowJo v10 software. Binding curves were generated by first gating on the live, HA-positive (*i.e.* HNV-G-positive) population. The GMFI (geometric mean fluorescence intensity) in the red channel of this gated, G-positive population quantified the level of ephrin-B ligand binding. The highest GMFI value obtained within each dilution series was normalized to 100%, and cell surface K_d_ values were calculated using GraphPad Prism.

### Pseudotyped virus

VSV particles pseudotyped with the HNV surface glycoproteins (HNVpp) were produced as previously described [26, 44, 54]. Briefly, particles were made from a VSV-ΔG-rLuc virus, a recombinant VSV derived from a full-length complementary DNA clone of the VSV Indiana serotype in which the VSV-G envelope protein is replaced by *Renilla* Luc. Pseudotyping was accomplished by transfecting HEK-293T cells (using BioT, BioLand Sci) with expression plasmids containing the codon-optimized C-terminally tagged F and G envelope glycoproteins of NiV (NiVpp), HeV (HeVpp), GhV (GhVpp), CedV (CedVpp), or the VSV-G glycoprotein itself (VSVpp), and then infecting with VSV-ΔG-rLuc (complemented with VSV-G). Pseudotype-containing media were clarified 48 h after infection, by centrifugation at 200 xg for 5 min. Supernatants were then loaded on a 20% sucrose cushion and subject to ultra-centrifugation for 3 h at 110,000 xg. Concentrated pseudoparticle pellets were then resuspended in Dulbecco’s Phosphate Buffered Saline (dPBS) and stored at −80 °C.

### Incorporation of henipaviral glycoproteins into HNVpp

Incorporation of F and G glycoproteins within NiVpp and CedVpp was determined by western blotting. Both F and G were detected using tag-specific anti-AU1 and anti-HA antibodies, respectively. Dilutions of NiVpp, CedVpp, and BALDpp (VSV virions bearing no glycoprotein) were lysed in 6× Laemmli buffer (5% β-mercaptoethanol final), boiled for 10 min, and separated on an Any kD Mini-PROTEAN TGX Precast Protein Gel before transfer to a PVDF membrane. Membranes were stained with a primary rabbit anti-HA pAb (Novus, catalogue number: NB600-363), diluted 1:2,000, and a primary rabbit anti-AU1 pAb (Novus, catalogue number: NB600-453), diluted 1:2,000. The membranes were then washed in PBS supplemented with 0.1% Tween-20 and incubated with Alexa 647-labelled goat anti-rabbit antibody (Thermo, catalogue number: A-21245), diluted 1:4,000. To control for loading, membranes were also stained for VSV matrix protein using a primary mouse anti-VSV-M mAb (Kerafast, catalogue number: EB0011), diluted 1:1,1000, and a secondary Alexa 647-labelled goat anti-mouse antibody (Thermo, catalogue number: A-21236), diluted 1:3,000. Membranes were imaged using a BioRad ChemiDoc and densitometry analysis was performed by using the BioRad Image Lab 5.1 software.

### Quantification of viral genome copies

Viral RNA was extracted from HNVpp preparations using the QIAamp viral RNA minikit (Qiagen), and subsequently reverse transcribed using the Tetro cDNA synthesis kit (Bioline). VSV genome copy number was quantified by qPCR, using the SensiFAST SYBR & Fluorescein kit (Bioline), utilizing genome-specific primers against the VSV Indiana L region (sequences available on request). Standard curves were generated by a serial dilution of the full-length VSV-ΔG-rLuc genomic plasmid.

### Quantification of viral entry

Target cells were grown in 96-well plates and infected with the pseudoviruses serially diluted in appropriate cell culture medium. For soluble ephrin-B entry inhibition experiments, the indicated amounts of soluble ephrin-B–Fc (R&D Systems) were incubated together with a pseudotyped virus for 1 h at 37 °C. The mixture of virus and soluble ephrin-B was then added to the target cells. For soluble envelope entry inhibition experiments, the indicated quantity of soluble HNV-G-Fc (production described above) or soluble Eph-B3-Fc (R&D Systems) were incubated with the adherent target cells for 1 h at 37 °C. Following incubation with soluble protein, pseudotyped virus was added. To assess viral entry into the wild-type and mutant ephrin expressing cell lines (Fig 6d), equivalent amounts of NiVpp and CedVpp (titrated to give roughly equivalent rLuc activities on CHO-B2 cells) were used to infect the distinct ephrin-expressing CHO-pgsA745 cell lines.

In all instances, the quantity of pseudotyped virus stock utilized was predetermined to fall within the linear dynamic range of *Renilla* Luc detection. Furthermore, in all HNVpp entry experiments, infected cells were washed with PBS and lysed 24 h post infection. Cell lysates were subsequently processed using a *Renilla* Luciferase Detection kit, according to the manufacturer’s directions (Promega). Luminescence intensity was measured using a Cytation3 Plate Reader.

### Quantification of ephrin-B mRNA transcripts in primary HUVECs

Ephrin-B1, -B2, and -B3 mRNA transcripts present in the primary HUVECs were quantified using qPCR. Total RNA was extracted from 3×10^5^ HUVECs with a NucleoSpin RNA isolation kit (Macherey-Nagel). Processed mRNA transcripts were reverse transcribed into cDNA using oligo(dT) primers and the Tetro cDNA synthesis kit (Bioline). qPCR was performed using the SensiFAST SYBR & Fluorescein kit (Bioline), using gene-specific primers (sequences available on request) for ephrin-B1, -B2, and -B3. Standard curves for each gene were generated by a serial dilution of the ephrin-B pcDNA3.1 expression plasmids. As a normalization control, hypoxanthine phosphoribosyltransferase (HPRT) copy numbers were also determined.

### HNV glycoprotein-mediated syncytia formation in U87 glioblastoma cells

Syncytia assays were performed by transfecting U87 cells with HNV-F and HNV-G expression plasmids, or an empty vector control (using Lipofectamine 2000, Thermo). Bright-field images were taken at 48 h post transfection with a Nikon Eclipse TE300 Diaphot Microscope. Images represent two random fields for each condition.

### Data deposition

The atomic coordinates and structure factors for CedV-G and CedV-G–ephrin-B1 will be deposited in the Protein Data Bank upon final acceptance of the manuscript.

## Acknowledgments

We are grateful to Diamond Light Source for beamtime (proposal MX19946), the staff of beamlines I02 and I24 for assistance with data collection, and to Helen M. Ginn for helpful discussions. We thank the MRC (MR/L009528/1 and MR/S007555/1 to T.A.B.), Academy of Finland (#309605 to I.R.), and NIH (NIAID R01 AI123449, R01 AI069317, and R21 AI115226 to B.L) for funding. K.A. was supported by the National Institute of Allergy and Infectious Diseases of the National Institutes of Health under Award Number F31-AI133943 & the Host-Pathogens Interactions Training Grant T32-AI007647-16 at the Icahn School of Medicine at Mount Sinai. B.L. also acknowledges the Ward Coleman estate for endowing the Ward-Coleman Chairs at the Icahn School of Medicine at Mount Sinai. The Wellcome Centre for Human Genetics is supported by Wellcome Centre grant 203141/Z/16Z. The Genotype-Tissue Expression (GTEx) Project was supported by the Common Fund of the Office of the Director of the National Institutes of Health, and by NCI, NHGRI, NHLBI, NIDA, NIMH, and NINDS. The data used for the analyses described in this manuscript (Supplementary Fig. 5) were obtained from the GTEx Portal dbGaP accession number <u>phs000424.v7.p</u>2 on 07/24/2019.

## Author contributions

R.P and K.A performed the experiments and analyzed results, alongside T.A.B and B.L. T.A.B and B.L devised and oversaw the study. I.R and K.H provided reagents and technical assistance. R.P, K.A, T.A.B, and B.L wrote the manuscript. R.P and K.A contributed equally.

## References

1. Hossain, M.J., et al., Clinical presentation of nipah virus infection in Bangladesh. Clin Infect Dis, 2008. 46(7): p. 977–84.

2. Luby, S.P., et al., Recurrent zoonotic transmission of Nipah virus into humans, Bangladesh, 2001-2007. Emerg Infect Dis, 2009. 15(8): p. 1229–35.

3. Arunkumar, G., et al., Outbreak Investigation of Nipah Virus Disease in Kerala, India, 2018. J Infect Dis, 2019. 219(12): p. 1867–1878.

4. WHO, 2018 Annual review of diseases prioritized under the Research and Development Blueprint 2018.

5. Gurley, E.S., et al., Person-to-person transmission of Nipah virus in a Bangladeshi community. Emerg Infect Dis, 2007. 13(7): p. 1031–7.

6. Ching, P.K., et al., Outbreak of henipavirus infection, Philippines, 2014. Emerg Infect Dis, 2015. 21(2): p. 328–31.

7. Nikolay, B., et al., Transmission of Nipah Virus – 14 Years of Investigations in Bangladesh. N Engl J Med, 2019. 380(19): p. 1804–1814.

8. Halpin, K., et al., Pteropid bats are confirmed as the reservoir hosts of henipaviruses: a comprehensive experimental study of virus transmission. Am J Trop Med Hyg, 2011. 85(5): p. 946–51.

9. Li, Y., et al., Antibodies to Nipah or Nipah-like viruses in bats, China. Emerg Infect Dis, 2008. 14(12): p. 1974–6.

10. Weiss, S., et al., Henipavirus-related sequences in fruit bat bushmeat, Republic of Congo. Emerg Infect Dis, 2012. 18(9): p. 1536–7.

11. Pernet, O., et al., Evidence for henipavirus spillover into human populations in Africa. Nat Commun, 2014. 5: p. 5342.

12. Drexler, J.F., et al., Bats host major mammalian paramyxoviruses. Nat Commun, 2012. 3: p. 796.

13. de Araujo, J., et al., Antibodies Against Henipa-Like Viruses in Brazilian Bats. Vector Borne Zoonotic Dis, 2017. 17(4): p. 271–274.

14. Mbu’u, C.M., et al., Henipaviruses at the Interface Between Bats, Livestock and Human Population in Africa. Vector Borne Zoonotic Dis, 2019. 19(7): p. 455–465.

15. Peel, A.J., et al., Continent-wide panmixia of an African fruit bat facilitates transmission of potentially zoonotic viruses. Nat Commun, 2013. 4: p. 2770.

16. Thibault, P.A., et al., Zoonotic Potential of Emerging Paramyxoviruses: Knowns and Unknowns. Adv Virus Res, 2017. 98: p. 1–55.

17. Wu, Z., et al., Novel Henipa-like virus, Mojiang Paramyxovirus, in rats, China, 2012. Emerg Infect Dis, 2014. 20(6): p. 1064–6.

18. Rissanen, I., et al., Idiosyncratic Mojiang virus attachment glycoprotein directs a host-cell entry pathway distinct from genetically related henipaviruses. Nat Commun, 2017. 8: p. 16060.

19. Marsh, G.A., et al., Cedar virus: a novel Henipavirus isolated from Australian bats. PLoS Pathog, 2012. 8(8): p. e1002836.

20. Schountz, T., et al., Differential Innate Immune Responses Elicited by Nipah Virus and Cedar Virus Correlate with Disparate In Vivo Pathogenesis in Hamsters. Viruses, 2019. 11(3).

21. Lee, B. and Z.A. Ataman, Modes of paramyxovirus fusion: a Henipavirus perspective. Trends Microbiol, 2011. 19(8): p. 389–99.

22. Bose, S., T.S. Jardetzky, and R.A. Lamb, Timing is everything: Fine-tuned molecular machines orchestrate paramyxovirus entry. Virology, 2015. 479-480: p. 518–31.

23. Plattet, P. and R.K. Plemper, Envelope protein dynamics in paramyxovirus entry. MBio, 2013. 4(4).

24. Pernet, O., Y.E. Wang, and B. Lee, Henipavirus receptor usage and tropism. Curr Top Microbiol Immunol, 2012. 359: p. 59–78.

25. El Najjar, F., A.P. Schmitt, and R.E. Dutch, Paramyxovirus glycoprotein incorporation, assembly and budding: a three way dance for infectious particle production. Viruses, 2014. 6(8): p. 3019–54.

26. Lee, B., et al., Molecular recognition of human ephrinB2 cell surface receptor by an emergent African henipavirus. Proc Natl Acad Sci U S A, 2015. 112(17): p. E2156–65.

27. Zeltina, A., T.A. Bowden, and B. Lee, Emerging Paramyxoviruses: Receptor Tropism and Zoonotic Potential. PLoS Pathog, 2016. 12(2): p. e1005390.

28. Bossart, K.N., et al., Functional studies of host-specific ephrin-B ligands as Henipavirus receptors. Virology, 2008. 372(2): p. 357–71.

29. Marsh, G.A. and L.-F. Wang, Hendra and Nipah viruses: why are they so deadly? Current Opinion in Virology, 2012. 2(3): p. 242–247.

30. de Wit, E. and V.J. Munster, Animal models of disease shed light on Nipah virus pathogenesis and transmission. J Pathol, 2015. 235(2): p. 196–205.

31. Park, A., et al., Nipah Virus C Protein Recruits Tsg101 to Promote the Efficient Release of Virus in an ESCRT-Dependent Pathway. PLoS Pathog, 2016. 12(5): p. e1005659.

32. Lo, M.K., et al., Distinct and overlapping roles of Nipah virus P gene products in modulating the human endothelial cell antiviral response. PLoS One, 2012. 7(10): p. e47790.

33. Mathieu, C., et al., Nonstructural Nipah virus C protein regulates both the early host proinflammatory response and viral virulence. J Virol, 2012. 86(19): p. 10766–75.

34. Basler, C.F., Nipah and hendra virus interactions with the innate immune system. Curr Top Microbiol Immunol, 2012. 359: p. 123–52.

35. Laing, E.D., et al., Rescue and characterization of recombinant cedar virus, a non-pathogenic Henipavirus species. Virol J, 2018. 15(1): p. 56.

36. Bowden, T.A., et al., Crystal structure and carbohydrate analysis of Nipah virus attachment glycoprotein: a template for antiviral and vaccine design. J Virol, 2008. 82(23): p. 11628–36.

37. Bowden, T.A., et al., Dimeric architecture of the Hendra virus attachment glycoprotein: evidence for a conserved mode of assembly. J Virol, 2010. 84(12): p. 6208–17.

38. Bowden, T.A., et al., Shared paramyxoviral glycoprotein architecture is adapted for diverse attachment strategies. Biochem Soc Trans, 2010. 38(5): p. 1349–55.

39. Yuan, P., et al., Structure of the Newcastle disease virus hemagglutinin-neuraminidase (HN) ectodomain reveals a four-helix bundle stalk. Proc Natl Acad Sci U S A, 2011. 108(36): p. 14920–5.

40. Welch, B.D., et al., Structure of the parainfluenza virus 5 (PIV5) hemagglutinin-neuraminidase (HN) ectodomain. PLoS Pathog, 2013. 9(8): p. e1003534.

41. Bradel-Tretheway, B.G., et al., Novel Functions of Hendra Virus G N-Glycans and Comparisons to Nipah Virus. J Virol, 2015. 89(14): p. 7235–47.

42. Pryce, R., et al., Structure-based classification defines the discrete conformational classes adopted by the arenaviral GP1. J Virol, 2018.

43. Negrete, O.A., et al., Two key residues in ephrinB3 are critical for its use as an alternative receptor for Nipah virus. PLoS Pathog, 2006. 2(2): p. e7.

44. Negrete, O.A., et al., Single amino acid changes in the Nipah and Hendra virus attachment glycoproteins distinguish ephrinB2 from ephrinB3 usage. J Virol, 2007. 81(19): p. 10804–14.

45. Bowden, T.A., et al., Structural basis of Nipah and Hendra virus attachment to their cell-surface receptor ephrin-B2. Nat Struct Mol Biol, 2008. 15(6): p. 567–72.

46. Xu, K., et al., Host cell recognition by the henipaviruses: crystal structures of the Nipah G attachment glycoprotein and its complex with ephrin-B3. Proc Natl Acad Sci U S A, 2008. 105(29): p. 9953–8.

47. Tamin, A., et al., Development of a neutralization assay for Nipah virus using pseudotype particles. J Virol Methods, 2009. 160(1-2): p. 1–6.

48. Chua, K.B., et al., Nipah virus: a recently emergent deadly paramyxovirus. Science, 2000.288(5470): p. 1432–5.

49. Wong, K.T., et al., Nipah virus infection: pathology and pathogenesis of an emerging paramyxoviral zoonosis. Am J Pathol, 2002. 161(6): p. 2153–67.

50. Erbar, S. and A. Maisner, Nipah virus infection and glycoprotein targeting in endothelial cells. Virol J, 2010. 7: p. 305.

51. Blits-Huizinga, C.T., et al., Ephrins and their receptors: binding versus biology. IUBMB Life, 2004. 56(5): p. 257–65.

52. Krissinel, E. and K. Henrick, Inference of macromolecular assemblies from crystalline state. J Mol Biol, 2007. 372(3): p. 774–97.

53. Nikolov, D.B., et al., Crystal Structure of the Ephrin-B1 Ectodomain: Implications for Receptor Recognition and Signaling. Biochemistry, 2005. 44(33): p. 10947–10953.

54. Negrete, O.A., et al., EphrinB2 is the entry receptor for Nipah virus, an emergent deadly paramyxovirus. Nature, 2005. 436(7049): p. 401–5.

55. Bonaparte, M.I., et al., Ephrin-B2 ligand is a functional receptor for Hendra virus and Nipah virus. Proc Natl Acad Sci U S A, 2005. 102(30): p. 10652–7.

56. Carithers, L.J., et al., A Novel Approach to High-Quality Postmortem Tissue Procurement: The GTEx Project. Biopreserv Biobank, 2015. 13(5): p. 311–9.

57. Lim, C.C., et al., Nipah viral encephalitis or Japanese encephalitis? MR findings in a new zoonotic disease. AJNR Am J Neuroradiol, 2000. 21(3): p. 455–61.

58. Ang, B.S.P., T.C.C. Lim, and L. Wang, Nipah Virus Infection. J Clin Microbiol, 2018. 56(6).

59. Liebl, D.J., et al., mRNA expression of ephrins and Eph receptor tyrosine kinases in the neonatal and adult mouse central nervous system. J Neurosci Res, 2003. 71(1): p. 7–22.

60. Yokoyama, N., et al., Forward signaling mediated by ephrin-B3 prevents contralateral corticospinal axons from recrossing the spinal cord midline. Neuron, 2001. 29(1): p. 85–97.

61. Kullander, K., et al., Ephrin-B3 is the midline barrier that prevents corticospinal tract axons from recrossing, allowing for unilateral motor control. Genes Dev, 2001. 15(7): p. 877–88.

62. Wong, K.T. and C.T. Tan, Clinical and pathological manifestations of human henipavirus infection. Curr Top Microbiol Immunol, 2012. 359: p. 95–104.

63. Aricescu, A.R., W. Lu, and E.Y. Jones, A time- and cost-efficient system for high-level protein production in mammalian cells. Acta Crystallogr D Biol Crystallogr, 2006. 62(Pt 10): p. 1243–50.

64. Chang, V.T., et al., Glycoprotein structural genomics: solving the glycosylation problem. Structure, 2007. 15(3): p. 267–73.

65. Walter, T.S., et al., A procedure for setting up high-throughput nanolitre crystallization experiments. Crystallization workflow for initial screening, automated storage, imaging and optimization. Acta Crystallographica Section D, 2005. 61(6): p. 651–657.

66. Newman, J., et al., Towards rationalization of crystallization screening for small-to medium-sized academic laboratories: the PACT/JCSG+ strategy. Acta Crystallogr D Biol Crystallogr, 2005. 61(Pt 10): p. 1426–31.

67. Winter, G., xia2: an expert system for macromolecular crystallography data reduction. Journal of Applied Crystallography, 2010. 43(1): p. 186–190.

68. McCoy, A.J., et al., Phaser crystallographic software. Journal of Applied Crystallography, 2007. 40(4): p. 658–674.

69. Emsley, P. and K. Cowtan, Coot: model-building tools for molecular graphics. Acta Crystallogr D Biol Crystallogr, 2004. 60(Pt 12 Pt 1): p. 2126–32.

70. Adams, P.D., et al., PHENIX: a comprehensive Python-based system for macromolecular structure solution. Acta Crystallogr D Biol Crystallogr, 2010. 66(Pt 2): p. 213–21.

71. Murshudov, G.N., A.A. Vagin, and E.J. Dodson, Refinement of macromolecular structures by the maximum-likelihood method. Acta Crystallogr D Biol Crystallogr, 1997. 53(Pt 3): p. 240–55.

72. Agirre, J., et al., Privateer: software for the conformational validation of carbohydrate structures. Nat Struct Mol Biol, 2015. 22(11): p. 833–4.

73. Lawrence, M.C., et al., Structure of the haemagglutinin-neuraminidase from human parainfluenza virus type III. J Mol Biol, 2004. 335(5): p. 1343–57.

74. Crennell, S., et al., Crystal structure of the multifunctional paramyxovirus hemagglutinin-neuraminidase. Nat Struct Biol, 2000. 7(11): p. 1068–74.

75. Kubota, M., et al., Trisaccharide containing alpha2,3-linked sialic acid is a receptor for mumps virus. Proc Natl Acad Sci U S A, 2016. 113(41): p. 11579–11584.

76. Stuart, D.I., et al., Crystal structure of cat muscle pyruvate kinase at a resolution of 2.6 A. J Mol Biol, 1979. 134(1): p. 109–42.

77. Felsenstein, J., PHYLIP – Phylogeny Inference Package (Version 3.2). Cladistics, 1989. 5: p. 164–166.

78. Corpet, F., Multiple sequence alignment with hierarchical clustering. Nucleic Acids Res, 1988. 16(22): p. 10881–90.

79. Gouet, P., X. Robert, and E. Courcelle, ESPript/ENDscript: Extracting and rendering sequence and 3D information from atomic structures of proteins. Nucleic Acids Res, 2003. 31(13): p. 3320–3.

