## Supplemental Figures for "A key region of molecular specificity orchestrates unique ephrin-B1 utilization by Cedar virus"

### Supplementary information

### Supplementary Table 1: Crystallographic data collection and refinement statistics

| **Data Collection** | **CedV-G** | **CedV-G–ephrin-B1** |
|---|---|---|
| Beamline | DLS I02 | DLS I24 |
| Wavelength (Å) | 0.9795 | 0.9686 |
| Space group | $P6_5$ | $P2_12_12_1$ |
| Cell dimensions | | |
| *a*, *b*, *c* (Å) | 201.5, 201.5, 112.9 | 112.8, 138.4, 235.4 |
| $\alpha$, β, $\gamma$ (°) | 90, 90, 120 | 90,90,90 |
| Resolution range (Å) | 58.16-2.78 (2.85-2.78) | 70.19-4.07 (4.14-4.07) |
| $R_{merge}$ | 0.164 (1.776) | 0.407 (2.192) |
| $R_{pim}$ | 0.037 (0.412) | 0.173 (0.914) |
| I/σ(I) | 15.4 (2.0) | 3.4 (0.9) |
| $CC_{1/2}$ | 0.998 (0.777) | 0.965 (0.359) |
| Completeness (%) | 99.9 (99.6) | 99.8 (92.4) |
| Multiplicity | 20.5 (19.5) | 6.5 (6.3) |
| Unique reflections | 65,603 (4,849) | 30,120 (1,373) |
| Wilson B-factor | 69.3 | 122.6 |
| **Refinement** | | |
| Resolution (Å) | 49.24-2.78 | 69.22-4.07 |
| No. reflections | 65,556 | 30,051 |
| $R_{work}/R_{free}$ | 0.201/0.227 | 0.276/0.34 |
| Protein chains in a.s.u | 2 | 10 |
| No. of atoms | 6,889 | 22,772 |
| Protein/ligand/water | 6,698/182/9 | 22,104/668/0 |
| *B*-factors Protein/ligand/water | 85.9/102.6/62 | 152.8/175.4/n.a |
| Ramachandran favored/allowed/outlier (%) | 93.9/6.0/0.1 | 92.7/7.2/0.1 |
| Root mean square deviations (RMSD) | | |
| Bond lengths (Å) | 0.003 | 0.004 |
| Bond angles (°) | 0.53 | 0.93 |

Values for the highest resolution shell are shown in parentheses.

**Supplementary Figure 1: Sequence alignment of ephrin-tropic HNV receptor-binding glycoproteins**

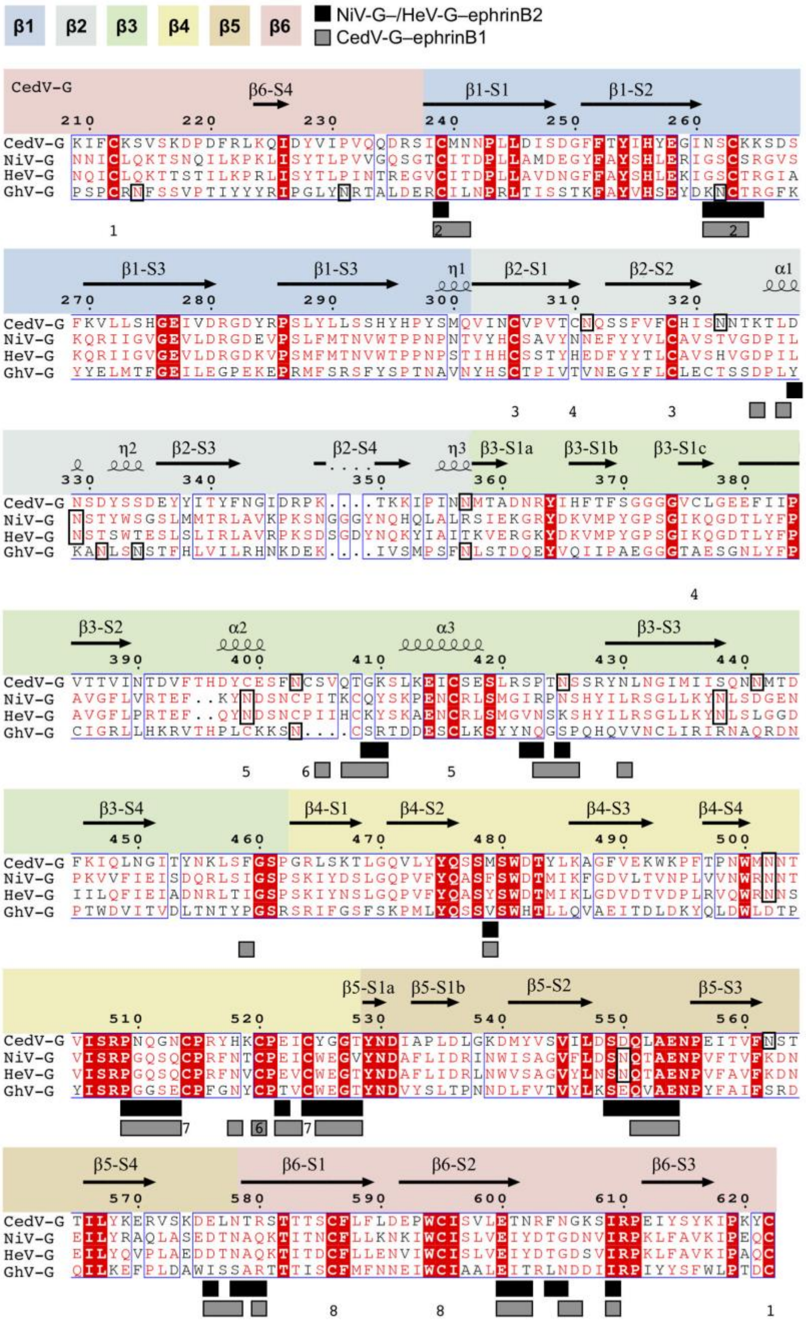


Sequence alignment of the β-propeller domains from CedV-G (NCBI reference sequence: YP_009094086.1), NiV-G (NP_112027.1), HeV-G (NP_047112.2), and GhV-G (YP_009091838.1). Absolutely conserved residues are shown as white on a red background, partially conserved residues are shown in red, and non-conserved residues are shown in black. Residues that interact with ephrin-B1 or ephrin-B2 are denoted with grey (ephrin-B1) or black (ephrin-B2) filled boxes beneath the sequence. Secondary structure elements of CedV-G are shown above the sequence and labelled according to type: β-strands are illustrated with arrows and labelled sequentially according to the β-blade (β1–β6) and strand number (S1–S4); α- and $3_{10}$ helices are illustrated as coils and labelled α1–α3 and η1–η3, respectively. Disulphide bonds within CedV-G are labelled beneath the sequence in order of appearance e.g '1' links cysteine residues 212 and 622. The putatively modified asparagine residues of N-linked glycosylation sequons are outlined with a black box. The blades of the β-propeller are colored in discrete sections, from blue to red. Sequence alignments were determined by MultAlin [78] and plotted using ESPript [79].

**Supplementary Figure 2: Electron density map features of N-linked glycans**

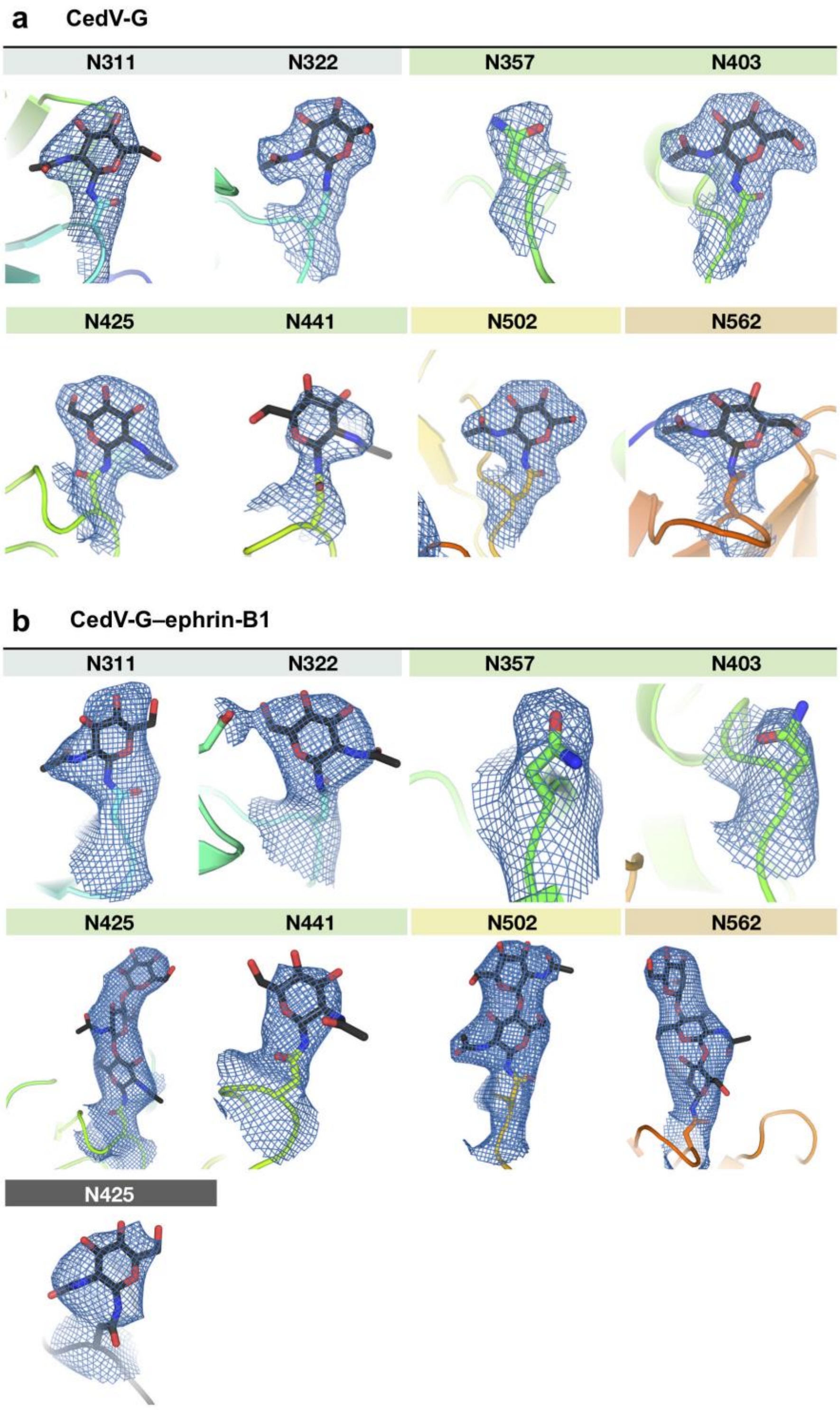


Electron density supported modelling of well-ordered N-linked glycans in the structures of CedV-G (**a**) and CedV-G–ephrin-B1 (**b**). The 2*Fo–Fc* electron density map (blue mesh; contoured to 1σ) are shown for all asparagine residues of N-linked glycosylation sequons and modelled asparagine-linked N-acetylglycosamine moieties, both shown as sticks. The main chain of CedV-G is displayed as a cartoon colored from blue to red (from N- to C-terminus) and ephrin-B1 is shown as a dark grey cartoon. Sticks are colored according to constituent elements (nitrogen is blue, oxygen is red, and carbon is colored according the moiety to which it belongs e.g grey if an ephrin-B1 atom). Glycans are labelled according to the sequence position of their associated asparagine residue.

**Supplementary Figure 3: CedV glycoproteins are efficiently incorporated into VSV particles but display reduced fusogenicity**

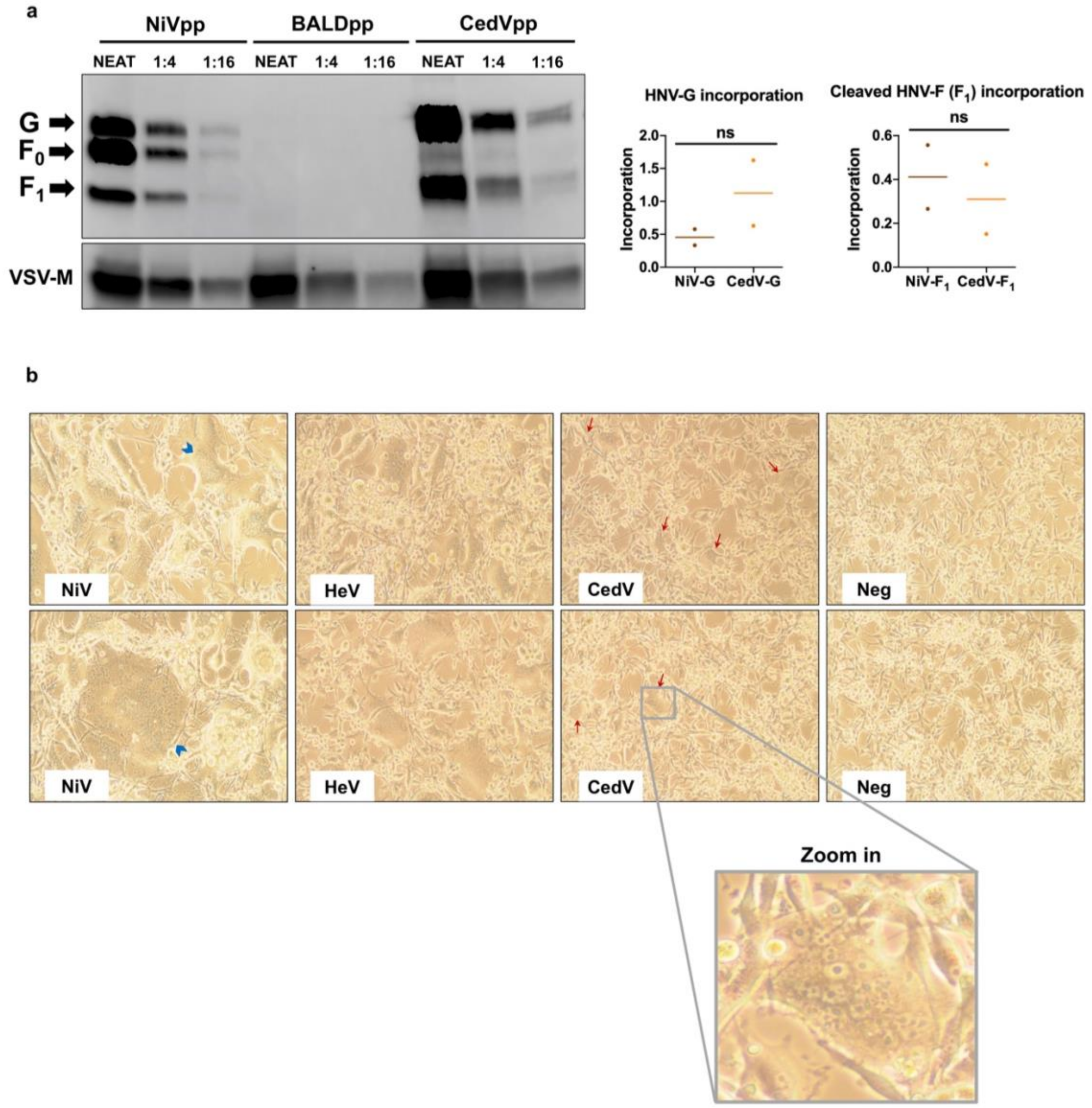


**a**, (*left*) Western blot analysis of the HNVpp. A 4-fold dilution series of NiVpp, CedVpp, and BALDpp was subjected to SDS-PAGE under reducing conditions. HNV-G and HNV-F proteins were detected using anti-HA and anti-AU1 antibodies, respectively. VSV matrix protein was used as a loading control and detected by a mouse anti-VSV-M monoclonal antibody. Arrows indicate the bands for HNV-G and both cleaved ($F_1$) and uncleaved ($F_0$) HNV-F. (*right*) Incorporation was quantified by western blotting densitometry. To calculate a level of incorporation, HNV-G and HNV-$F_1$ band intensity was normalized to that of VSV matrix. Data shown are the individual normalized densitometry values for two dilution points within the linear dynamic range in the 4-fold series. Horizontal dashes represent the mean from the two points. Statistical significance for the indicated comparisons were evaluated using a two-tailed unpaired t-test, n/s denotes no significance. **b**, Co-expression of CedV-F and CedV-G induced syncytia formation in U87 cells that were smaller than syncytia formed by NiV-F/-G and HeV-F/-G. CedV-F/G, NiV-F/G, HeV-F/G, or empty vector control were transfected into U87 glioblastoma cells. Images were taken 48 h post transfection. Blue arrowheads highlight large syncytia from NiV glycoprotein-mediated fusion and red arrows highlight the reduced syncytia generated from CedV glycoprotein-mediated fusion.

**Supplementary Figure 4: Electron density map features at a key receptor interaction site**

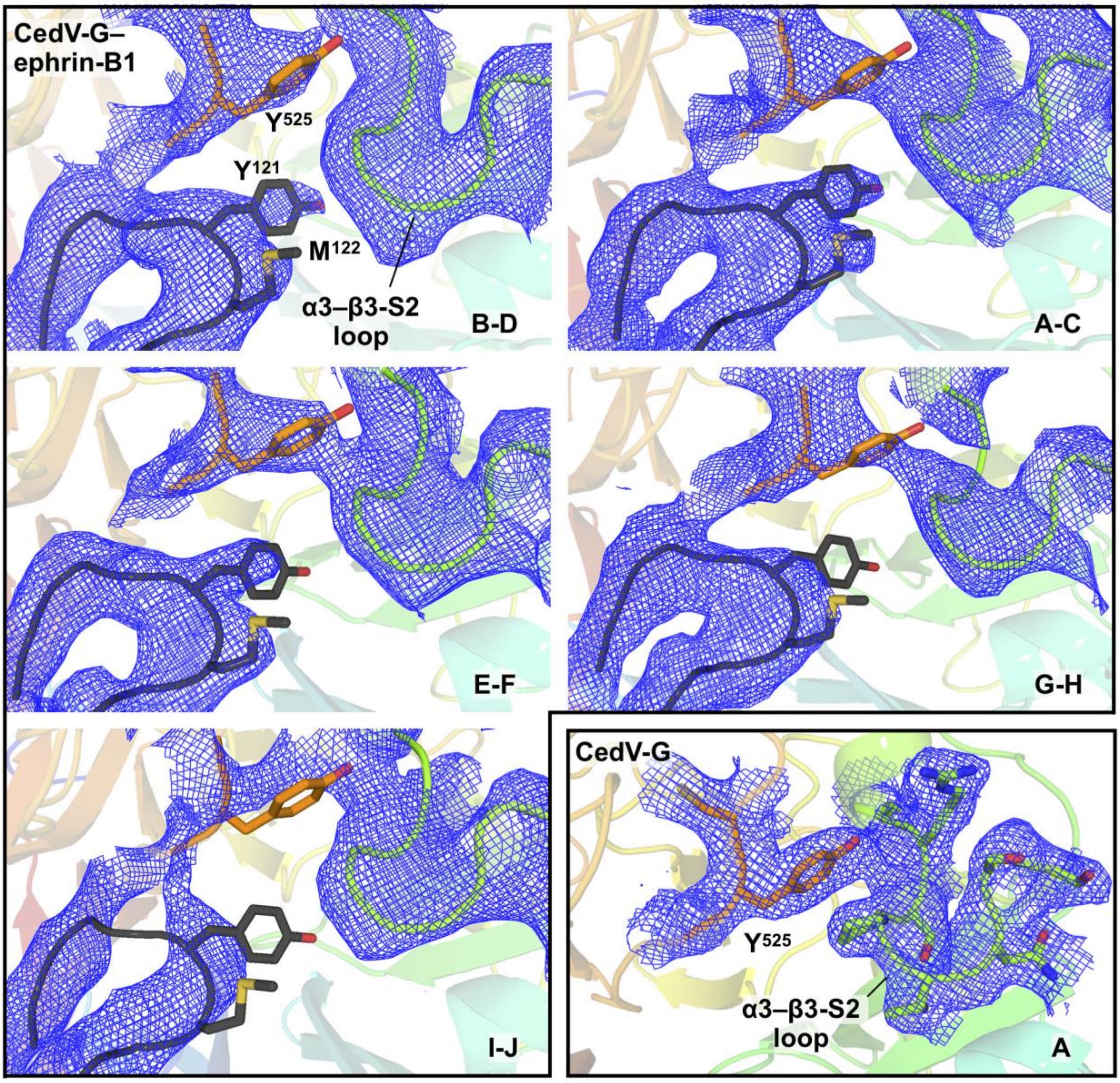


2*Fo*–*Fc* electron density maps (blue mesh; contoured to 1σ) are shown for a key region at the CedV-G–ephrin-B1 interface. The side chains of ephrin-B1 residues comprising the YM motif (Y121 and M122), and the CedV-G residue Y525, are shown as sticks and colored according to constituent elements. The α3–β3-S2 loop (residues 420–427) is shown as a cartoon in the CedV-G–ephrin-B1 structure with the addition of side chains in unliganded CedV-G. CedV-G is colored in a gradient from N- to C-terminus (blue to red) and ephrin-B1 is shown in dark grey. The a.s.u. chains to which each panel correspond is indicated in the bottom right. The complex comprising chains I and J exhibited regions of considerably poorer quality electron density than other a.s.u. molecules. Assessment of complexes B-D, A-C, and E-F, and the unliganded CedV-G structure (bottom right panel) enabled confidence in model building and allowed the formulation of structure-guided hypotheses that were tested with functional assays.

**Supplementary Figure 5: RNA-seq expression data of EFNB1, EFNB2, and EFNB3 in select human tissues**

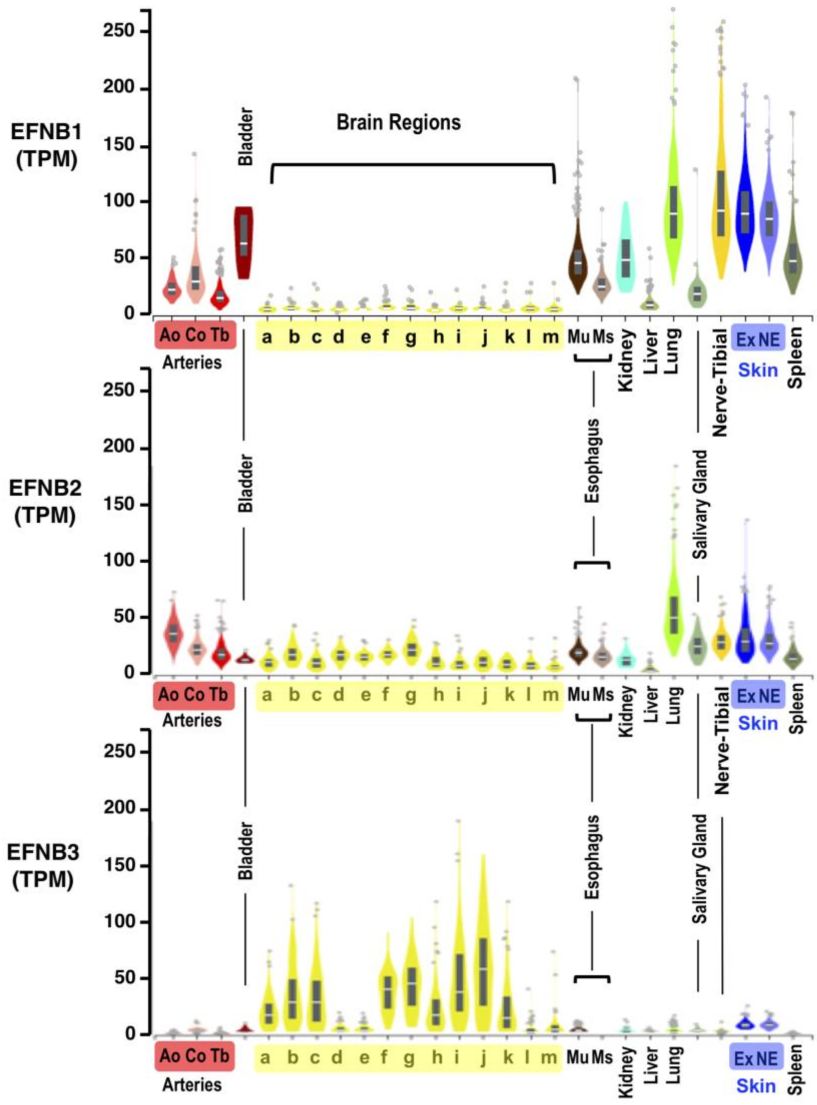


Expression values are shown in TPM (Transcripts Per Million), calculated from a gene model with isoforms collapsed to a single gene. Box plots are shown as median and 25$^{th}$ and 75$^{th}$ percentiles; points are displayed as outliers if they are above or below 1.5 times the interquartile range. Data was obtained through the Genotype-Tissue Expression Project (GTExPortal.org) where samples were collected from over 50 non-disease tissue sites from close to 1,000 individuals. Data for ephrin-B1 (EFNB1, top panel), ephrin-B2 (EFNB2, middle panel), and ephrin-B3 (EFNB3, bottom panel) are shown for the selected tissues listed alphabetically: **Arteries** (Ao, Aorta; Co; Coronary; Tb, Tibial) | **Bladder** | **Brain Regions** (**a** = Amygdala, **b** = Anterior cingulate cortex (BA24), **c** = Caudate (basal ganglia), **d** = Cerebellar hemisphere, **e** = Cerebellum, **f** = Cortex, **g** = Frontal Cortex (BA9), **h** = Hippocampus, **i** = Hypothalamus, **j** = Nucleus accumbens (basal ganglia), **k** = Putamen (basal ganglia), **l** = Spinal cord (cervical c-1), **m** = Substantia nigra) | **Esophagus** (Mu = Mucosa, Ms = Muscularis) | **Kidney** | **Liver** | **Lung** | **Salivary Gland** | **Nerve-Tibial** | **Skin** (Ex = Sun exposed; NE = Sun not exposed) | **Spleen**.
